# Subthalamic nucleus stabilizes movements by reducing neural spike variability in monkey basal ganglia: chemogenetic study

**DOI:** 10.1101/2021.04.13.439559

**Authors:** Taku Hasegawa, Satomi Chiken, Kenta Kobayashi, Atsushi Nambu

## Abstract

The subthalamic nucleus (STN) projects to the external pallidum (GPe) and internal pallidum (GPi), the relay and output nuclei of the basal ganglia (BG), respectively, and plays an indispensable role in controlling voluntary movements. To elucidate the neural mechanism by which the STN controls GPe/GPi activity and movements, we utilized a chemogenetic method to reversibly suppress the motor subregion of the STN in three macaque monkeys (*Macaca fuscata*, both sexes) engaged in reaching tasks. Systemic administration of chemogenetic ligands prolonged movement time and increased spike train variability in the GPe/GPi, but only slightly affected firing rate modulations. Across-trial analyses revealed that the irregular discharge activity in the GPe/GPi coincided with prolonged movement time. STN suppression also induced excessive abnormal movements in the contralateral forelimbs, which was preceded by STN and GPe/GPi phasic activity changes. Our results suggest that the STN stabilizes spike trains in the BG and achieves stable movements.

## Introduction

The subthalamic nucleus (STN) is small, but it occupies an important position in the basal ganglia (BG) circuitry. The STN receives cortical inputs directly through the cortico-STN pathway and indirectly through the cortico-striato-external pallido (GPe)-STN pathways^1^ and sends a glutamatergic projection to the GPe and internal pallidum (GPi). The GPe innervates all other nuclei in the BG^2,3^, whereas the GPi is the output nucleus of the BG^4,5^. Therefore, the STN affects the activity of all BG nuclei as well as the output nucleus.

The STN also plays pivotal roles in normal functions and disease conditions of the BG. Lesions or chemical blockade of the STN induces involuntary movements known as hemiballism^6–8^. Abnormal activity of STN neurons has been reported in Parkinson’s disease (PD)^9–12^, and it is suggested that the reciprocal excitatory and inhibitory connection between the STN and GPe is the source of pathologic oscillation associated with PD^13,14^. Moreover, lesions or deep brain stimulation (DBS) in the STN can ameliorate the motor symptoms of PD^15–17^.

These clinical effects are consistent with the classical BG model; the cortico-striato-GPi *direct* pathway facilitates movements, whereas the cortico-STN-GPi *hyperdirect* and cortico-striato-GPe-STN-GPi *indirect* pathways suppress movements^5,8,18,19^. This model hypothesizes that the STN inhibits and/or cancels movements, which is supported by human neuroimaging and electrophysiologic recording/stimulation studies of the STN^20–22^. However, the activity of STN neurons changes in relation to simple limb or eye movements as well^23,24^, and such movement-related activity is not easily explained from the perspective of movement suppression. It has been argued that the STN activates antagonist muscles necessary for stopping movements, or that it suppresses competing motor programs, thus allowing the *direct* pathway to release only a selected motor program^18,19,25^. Furthermore, pharmacologic activation of the STN induces involuntary movements on the contralateral side^26,27^. These previous observations suggest that the STN endows the BG circuitry with more complex neural computations than the simple dichotomy of movement facilitation and suppression.

To clarify the functional role of the STN in motor control, in the present study, we utilized the Designer Receptors Exclusively Activated by a Designer Drug (DREADD) technology to manipulate the neural activity in the STN of macaque monkeys. Although DREADDs have been utilized widely in rodents, only a few applications in non-human primates are reported, and electrophysiologic evaluation at the single-neuron level is lacking. Here, we showed that administration of DREADD ligands mildly suppressed STN activity, which was sufficient to induce abnormal involuntary movements and extend the movement time. Single-unit recordings revealed that pauses and spike train variability increased in both GPe and GPi neurons, whereas their movement-related activity was slightly affected. Our findings thus suggest a novel role for the STN in stabilizing spike trains in the GPe/GPi to achieve stable motor control.

## Results

### Mild suppression of STN activity using DREADD

The STN motor subregion involved in control of the forelimbs was identified based on neuronal responses to electrical stimulation of the forelimb regions of the primary motor cortex (M1) and supplementary motor area (SMA), i.e., biphasic early and late excitation via the cortico-STN *hyperdirect* and cortico-striato-GPe-STN *indirect* pathways, respectively (Fig. 1a, magenta and green arrows)^1,28,29^. An adeno-associated virus (AAV) vector that co-expresses an inhibitory DREADD receptor, hM4Di, and enhanced green fluorescent protein (EGFP) was injected unilaterally into the identified STN motor subregion. Histologic examination of monkeys E and K revealed EGFP expression in the dorsolateral part of the posterior STN (Fig. 1b and Supplementary Fig. 1), corresponding to the motor subregion^24,29–31^.

**Fig. 1.**
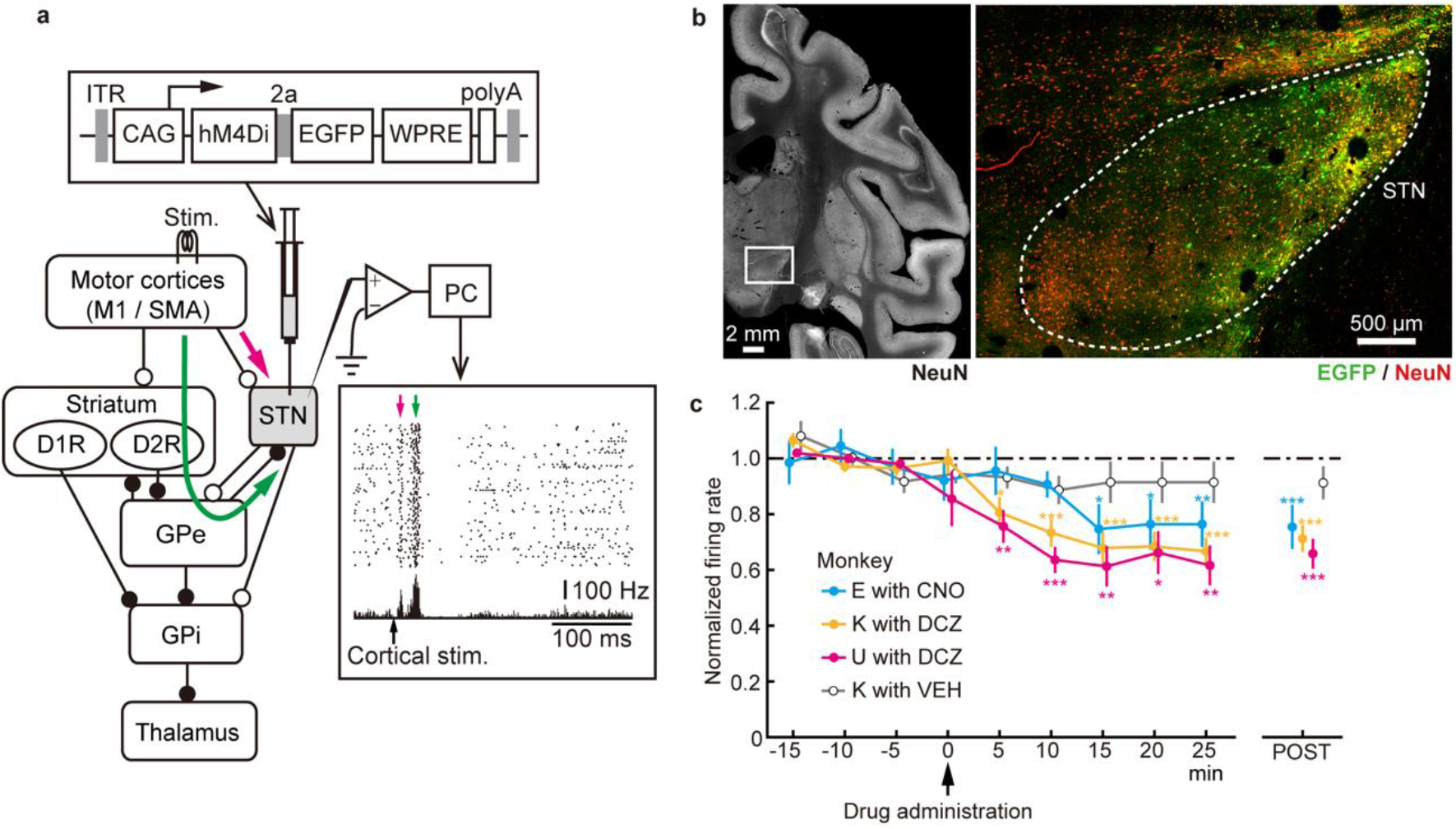
Suppression of STN activity using DREADD. **a**, Experimental overview. The motor subregion of the STN was identified by the characteristic biphasic excitation (bottom right inset) induced by electrical stimulation of the motor cortices, followed by infusion of an AAV vector co-expressing hM4Di and EGFP (top inset). Magenta and green arrows indicate early and late excitation. **b**, Histologic confirmation of AAV transduction with anti-GFP (green) and NeuN (red) antibodies in monkey K. The brain region indicated by an open box in the left image is enlarged on the right. **c**, Effects of systemic administration of CNO (1.0 mg/kg, i.v.), DCZ (0.1 mg/kg, i.m.), or vehicle (VEH) on baseline firing rates of STN neurons. STN activity was normalized based on activity during the PRE period (from −15 to 0 min) and averaged in 5-min bins (left) and in the POST periods (right; CNO, from 15 to 25 min; DCZ, from 10 to 25 min). Error bars indicate SEM. * P < 0.05, ** P < 0.01, *** P < 0.001, one-tailed Wilcoxon signed rank test with Bonferroni correction (n = 12, 24, and 20 neurons for monkeys E, K, and U, respectively).

After receptor expression (>3 weeks), neuronal recordings and behavioral observations were initiated. A ligand for hM4Di, clozapine N-oxide (CNO; 1 mg/kg, i.v., to monkey E) or newly developed deschloroclozapine (DCZ; 0.1 mg/kg, i.m., to monkeys K and U)^32^, was administered systemically. The efficacy of chemogenetic suppression was determined by recording unit activity in the STN. The baseline firing rates of the STN neurons began to decrease at 15 min after administration of CNO and 5 min after DCZ administration (Fig. 1c), consistent with the rapid delivery of DCZ to the brain^32^. The firing rates decreased to 65-75% of the rates before ligand administration (monkey E, 75.4 ± 7.9%, P < 0.001, n = 12; monkey K, 71.2 ± 4.8%, P < 0.001, n = 24; monkey U, 65.8 ± 5.4%, P < 0.001, n = 20; Wilcoxon signed rank test), whereas administration of vehicle had no effect (monkey K, 91.2 ± 7.9%, P = 0.1, n = 15).

Lesions or chemical blockade of the STN induces involuntary movements known as hemiballism^6–8^. In the present study, administration of DREADD ligands induced involuntary movements in all three monkeys, i.e., irregular repetitive movements in the contralateral arms without clear purpose while the monkeys sat quietly in a chair without performing any task (Supplementary Video 1; monkey E; occurrence rate, 0.77 min^−1^; duration, 19.0 ± 16.9 s, mean ± SD). No abnormal movements were noted with other body parts, such as the eyes, hindlimbs, or contralateral forelimb. The involuntary movements began approximately 20 min (CNO) or 5 min (DCZ) after ligand administration and continued for >2 h. Involuntary movements following ligand administration were observed repeatedly throughout the experimental periods (58, 85, and 75 weeks after AAV vector injection in monkeys E, K, and U, respectively). These histologic, electrophysiologic, and behavioral observations indicated that the forelimb motor subregion of the STN was successfully targeted and that DREADD expression was stable for >1 year.

### Movements disturbed by STN suppression

Each monkey was trained to perform a reach-and-pull task composed of externally triggered (ET) and self-initiated (SI) trials using its contralateral hand (Fig. 2a); the monkey was required to initiate reaching either immediately after ‘Go’ cue presentation in ET trials or with a delay of 1.5 s without any explicit Go cue in SI trials. ET and SI trials were randomly presented, and the trial type was indicated by the color of an LED (Task cue): blue for ET trials and red for SI trials. In ET trials, the blue LED turned green (Go cue) with a delay of 1-2 s, and then the monkey must release the home lever within 0.5 s and pull the front lever to be rewarded. In SI trials, the monkey must wait and maintain the home position for >1.5 s after Task cue and then initiate reaching movements.

**Fig. 2.**
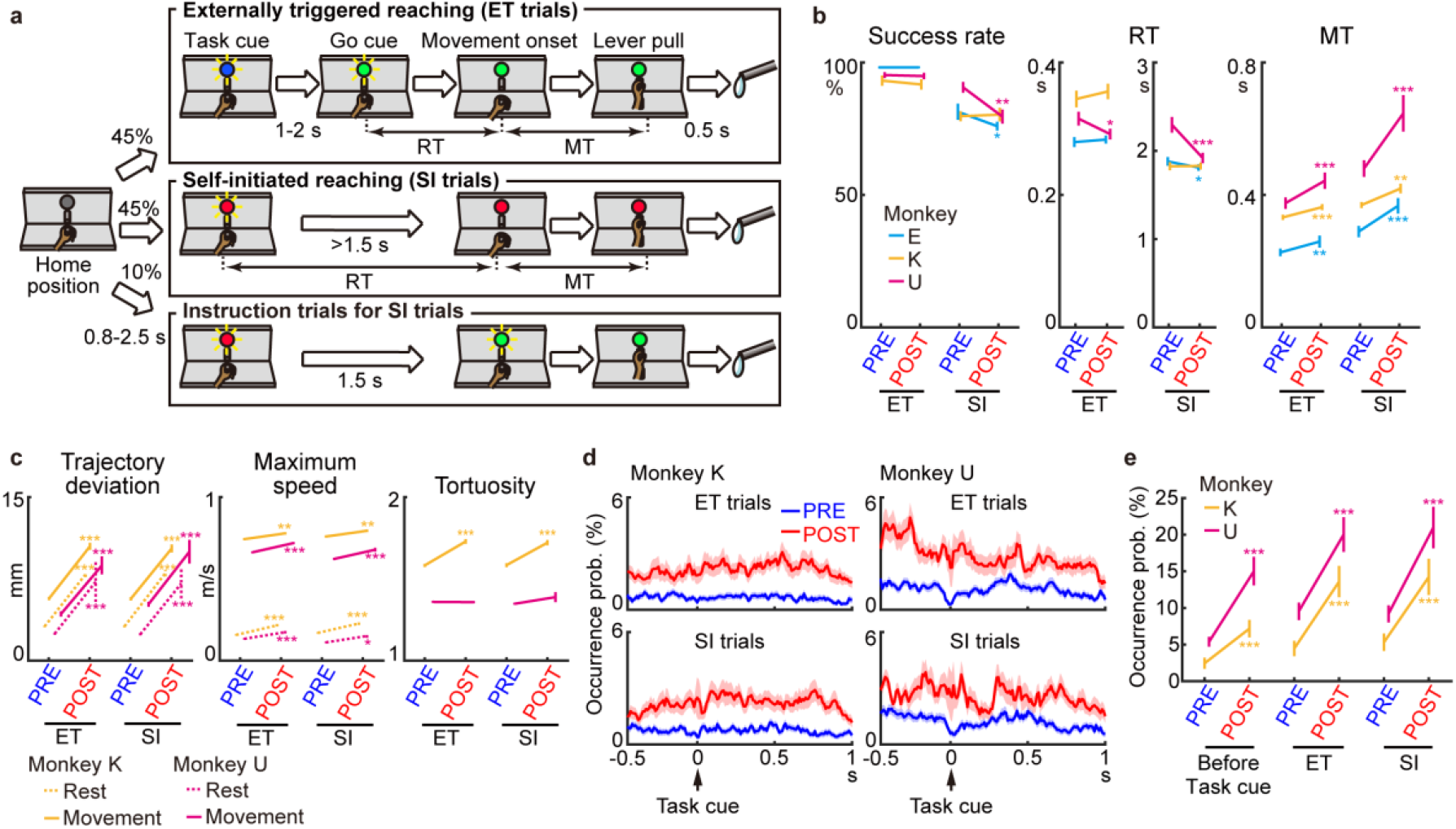
Motor effects of STN suppression on performance of a reach-and-pull task. a, Custom reach-and-pull task. See Online Methods for details. RT, reaction time; MT, movement time. **b**, Success rate, RT, and MT during the task using the hand contralateral to the AAV injection side before (PRE) and after (POST) the administration of a DREADD ligand. Error bars indicate SEM. * P < 0.05, ** P < 0.01, *** P < 0.001, two-tailed Wilcoxon signed rank test (n = 14, 15, and 20 sessions for monkeys E, K, and U, respectively). **c**, Analyses of wrist trajectories for monkeys K and U. See Supplementary Fig. 3. Trajectory deviation and maximum speed were calculated during the period −1 to 0 s relative to Movement onset (Rest), 0 to 1 s (Movement), and tortuosity during the period −0.5 to 0.5 s. Error bars indicate SEM. * P < 0.05, ** P < 0.01, *** P < 0.001, two-tailed Mann-Whitney *U* test (monkey K, 179 ET and 155 SI trials; monkey U, 101 ET and 103 SI trials). **d**, Occurrence probability of involuntary movements detected as fluctuations of the home lever (n = 44 and 43 sessions for monkeys K and U, respectively). Bin width, 1 ms. Shading indicates SEM. **e**, Occurrence probability of involuntary movements during the 0.5 s preceding (before Task cue) and 1 s following Task cue (ET and SI). Error bars indicate SEM. *** P < 0.001, two-tailed Wilcoxon signed rank test.

After CNO or DCZ administration, reaching motions became unstable, and the monkeys required more time to grab the front lever in some ET and SI trials, as observed by an increase in movement time (MT) in all three monkeys (Fig. 2b *right*; P < 0.01 for all monkeys in both ET and SI trials; Wilcoxon signed rank test; n = 14, 15, and 20 for monkeys E, K, and U, respectively). The success rate (Fig. 2b *left*; monkey E, P < 0.05; monkey U, P < 0.01) and reaction time (RT; Fig. 2b *middle*; monkey E, P < 0.05; monkey U, P < 0.01) of monkeys E and U decreased in SI trials. There was no change in the success rate and RT in the ET trials, except for a decrease in RT of monkey U (Fig. 2b *middle*; P < 0.05). No significant effects were observed on the success rate, RT, or MT after vehicle administration (Supplementary Fig. 2a; P > 0.05 for all monkeys in both ET and SI trials; Wilcoxon signed rank test; n = 8, 12, and 10 for monkeys E, K and U, respectively), or in task performance using the ipsilateral hand (Supplementary Fig. 2b; P > 0.05 for all monkeys in both ET and SI trials; n = 8, 9, and 9), indicating that any off-target effects of the DREADD ligands or their effects on STN non-motor functions were minimal.

Three-dimensional (3D) trajectories of the contralateral shoulder, elbow, wrist, and hand during reaching movements were reconstructed from RGB (x-y) and depth (z) images in monkeys K and U (Supplementary Fig. 3), and trajectories of the wrist position were statistically analyzed (Fig. 2c). The trajectory deviation increased in both monkeys (Fig. 2c *left* Movement): monkey K (ET trials, P < 10^−39^, n = 179; SI trials, P < 10^−28^, n = 155; Mann-Whitney *U* test) and monkey U (ET, P < 10^−14^, n = 101; SI, P < 10^−8^, n = 103). The maximum speed increased in both monkeys (Fig. 2c *middle* Movement): monkey K (ET, P < 0.01; SI, P < 0.01) and monkey U (ET, P < 10^−8^; SI, P < 10^−5^). The trajectory tortuosity, a measure of the bending and curving of a path, increased in monkey K (Fig. 2c *right*; ET, P < 10^−16^; SI, P < 10^−9^) but not monkey U (ET, P = 0.6; SI, P = 0.5). These results indicate that the longer MT in the POST period (Fig. 2b *right*) was due to high variability in reaching movements, rather than slow movements. In addition, analysis of the trajectory during the delay period (i.e., at rest) revealed task-irrelevant involuntary movements. The trajectory deviation increased in both monkeys (Fig. 2c *left* Rest): monkey K (ET, P < 10^−47^; SI, P < 10^−40^) and monkey U (ET, P < 10^−12^; SI, P < 10^−13^). In addition, the maximum speed increased in both monkeys (Fig. 2c *middle* Rest): monkey K (ET, P < 10^−7^; SI, P < 10^−5^) and monkey U (ET, P < 10^−4^; SI, P < 0.05). These changes were not observed in either monkey after vehicle administration: monkey K (Supplementary Fig. 2c; P > 0.05 for all measures and conditions; n = 74 ET and 70 SI trials) and monkey U (P > 0.05; n = 45 ET and 54 SI trials). Thus, excessive movements occurred both during movements and at rest, resulting in disturbed reaching and involuntary movements, respectively.

**Fig. 3.**
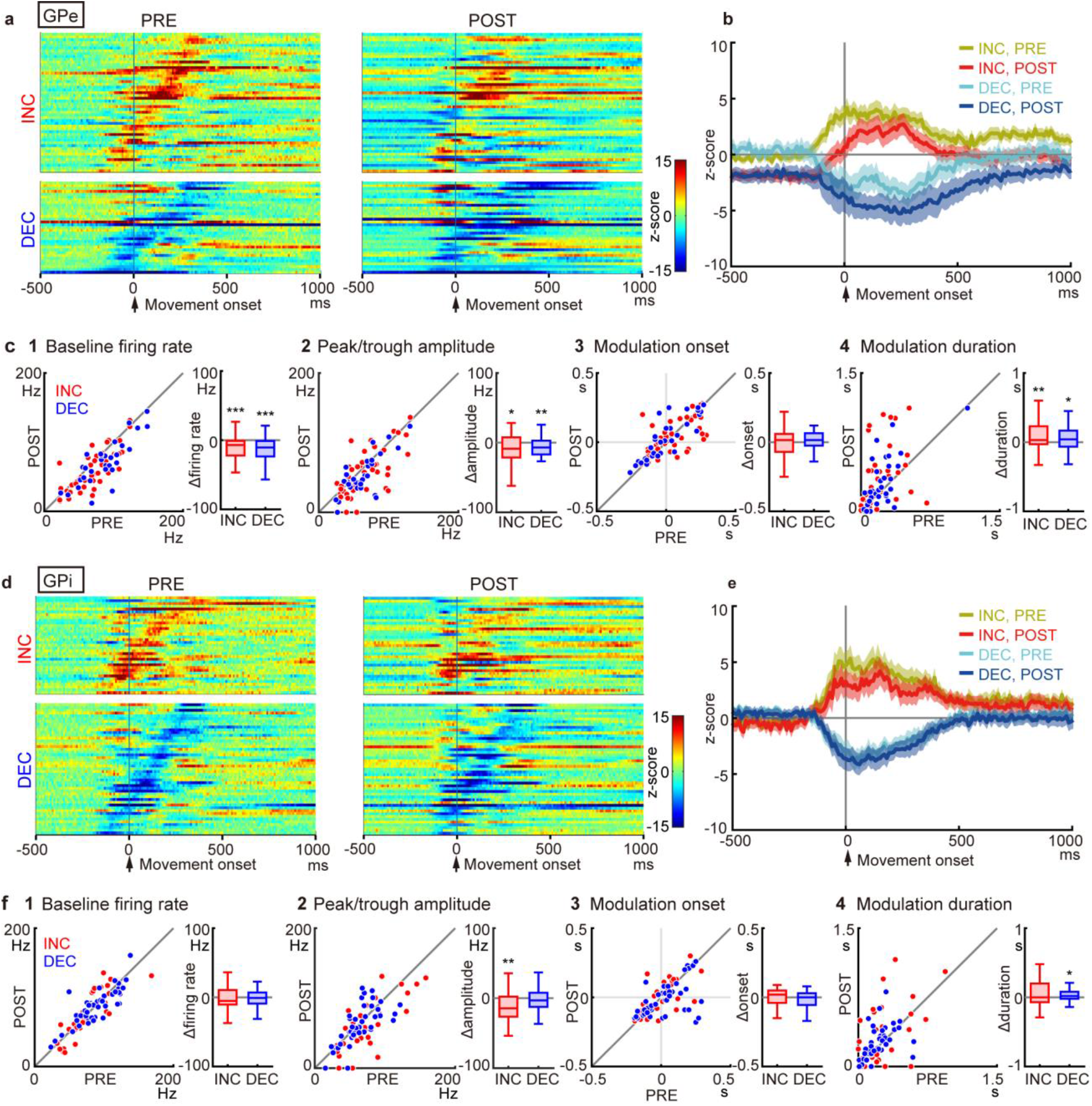
Changes in movement-related activity of GPe/GPi neurons after STN suppression. a, Heat maps for GPe neurons (n = 79) classified as INC or DEC types. Firing rates were converted to z-scores using the baseline during the 500 ms preceding Task cue in the PRE period. Neurons are sorted by the onset of movement-related activity in the PRE period (left). The activity of the same neuron in the POST period is shown on the same row (right). Bin width, 10 ms. **b**, Population-averaged PETHs of INC- and DEC-type GPe neurons in the PRE and POST periods. Solid lines and shading indicate mean and SEM, respectively. **c**, Change in PETHs between the PRE and POST periods. **c1**, baseline firing rate, calculated during the 500 ms preceding Task cue; **c2**, peak (INC) or trough (DEC) amplitude of the PETHs; **c3**, onset of movement-related modulations; **c4**, duration of movement-related modulations. In each box plot, an inner horizontal line indicates median; box, interquartile range (25^th^ and 75^th^ percentiles); whiskers, maximum and minimum values within 1.5 times the interquartile range from the upper and lower quartiles. **d-f**, Same as **(a-c)** but for GPi neurons (n = 78). Error bars indicate SEM. * P < 0.05, ** P < 0.01, *** P < 0.001, two-tailed Wilcoxon signed rank test.

Involuntary movements were also detected as fluctuations of the home lever position at rest in monkeys K and U. In both monkeys, involuntary movements became more frequent in the POST period before and after Task cue in both ET and SI trials (Fig. 2d). A significant increase in the occurrence of fluctuations was observed before Task cue (Fig. 2e; monkey K, from 2.5% to 7.2%, P < 10^−4^, n = 44; monkey U, from 5.4% to 15.0%, P < 10^−4^; Wilcoxon signed rank test), after Task cue in ET trials (monkey K, from 4.5% to 13.6%, P < 10^−5^; monkey U, from 9.5% to 20.0%, P < 10^−4^), and after Task cue in SI trials (monkey K, from 5.3% to 14.2%, P < 10^−5^; monkey U, from 9.2% to 21.0%, P < 0.001).

Electromyogram (EMG) of the biceps brachii and triceps brachii muscles of monkeys K and U was obtained during the task (Supplementary Fig. 4). Task-irrelevant phasic EMG activity was observed, corresponding to involuntary movements. In the POST period, abnormal EMG activity was induced more frequently after Task cue, and the movement-related EMG activity tended to increase, consistent with the abovementioned trajectory analyses.

**Fig. 4.**
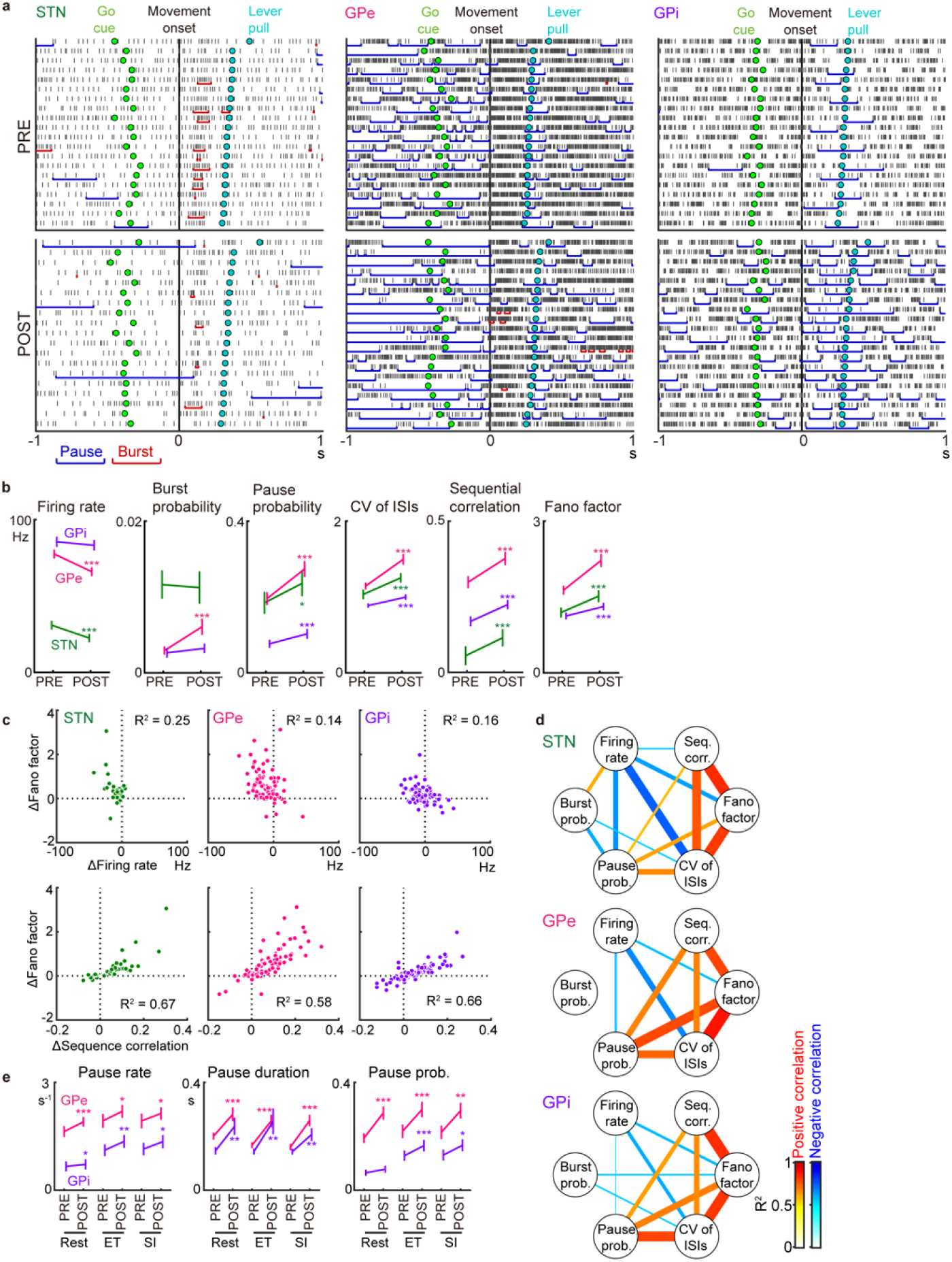
Firing pattern changes in the STN/GPe/GPi after STN suppression. a, Examples of STN/GPe/GPi neuronal activity in ET trials in the PRE and POST periods of STN suppression. Spikes are aligned with Movement onset, and trials are sorted by MTs. Vertical lines represent spikes; horizontal red and blue lines indicate bursts and pauses, respectively. **b**, Statistical analyses of spike trains. Firing rate, burst and pause probabilities, CV of ISIs, sequential correlation, and the Fano factor were calculated from −1 to 1 s relative to Movement onset in both the ET and SI trials. Error bars indicate SEM. * P < 0.05, ** P < 0.01, *** P < 0.001, two-tailed Wilcoxon signed rank test (n = 44 STN, 79 GPe, and 78 GPi neurons). **c**, Correlation between changes in the Fano factor and changes in firing rate (above) and sequential correlation (below). Each dot represents a change in a firing property of each neuron (POST – PRE). *R*^*2*^, squared Pearson correlation. **d**, Network representation for all pairwise correlations between the firing properties in (**b**). Nodes and links represent spike parameters and significant correlations, respectively. Line width and color intensity indicate the strength of positive (red) and negative (blue) correlations. **e**, Pauses in the GPe/GPi during the PRE and POST periods. Pause rate, pause duration, and pause probability were analyzed during separate task periods: from −0.5 to 0 s relative to Task cue (Rest) and from -0.2 to 0.3 s (ET and SI) relative to Movement onset.

### Diminished cortically evoked responses in GPe/GPi neurons by STN suppression

Single-unit GPe/GPi activity was recorded using a 16-channel linear electrode in 78 sessions. Of 198 neurons (101 GPe and 97 GPi) examined, 165 (83 GPe and 82 GPi) responded to M1 and/or SMA stimulation and were classified as ‘high-frequency discharge and pause’ (HFD-P) GPe, ‘low-frequency discharge and burst’ (LFD-B) GPe, ‘high-frequency discharge’ (HFD) GPi, or ‘low-frequency discharge’ neurons based on firing rates and patterns (Table 1)^33^. HFD-P GPe neurons (79/83) and HFD GPi neurons (78/82) were major and further analyzed, whereas LFD-B GPe neurons (4/83) are shown separately (Supplementary Fig. 5). STN activity was also recorded in another 32 sessions. Of 59 STN neurons examined, 44 responded to M1 and/or SMA stimulation (Table 1).

**Table 1.**
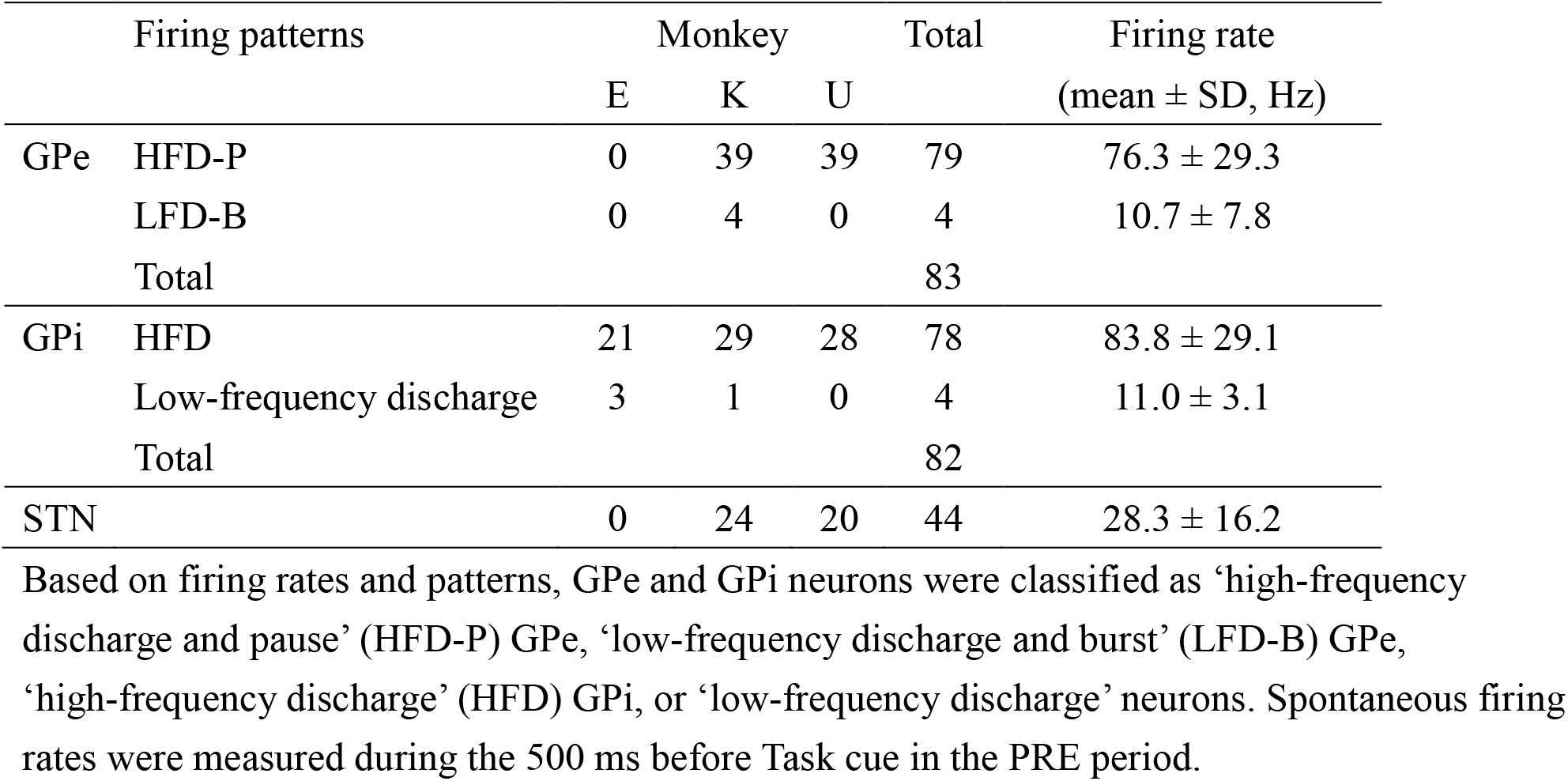
Number of STN, GPe, and GPi neurons recorded.

**Fig. 5.**
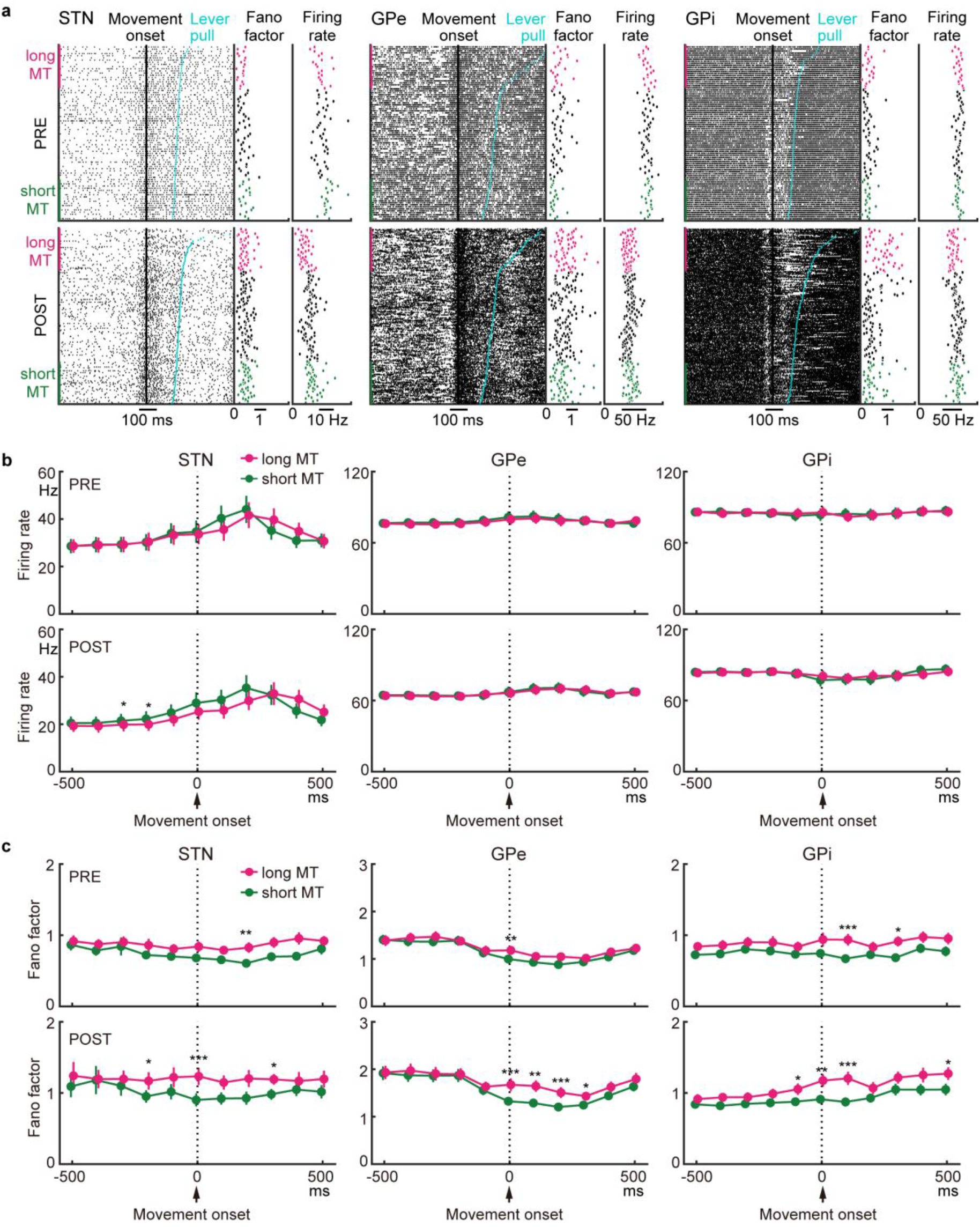
Neural activity correlated with disturbance in reaching movements after STN suppression. a, Examples of STN/GPe/GPi neuronal activity with the Fano factor and firing rate calculated in each trial. Spikes are aligned with Movement onset, and trials are sorted by MTs. In each neuron, the short- and long-MT trials were defined as trials with MTs below the 25^th^ and above the 75^th^ percentile, respectively. Both ET and SI trials were combined. **b**, Population-averaged firing rates of STN/GPe/GPi neurons for the short- and long-MT trials in the PRE and POST periods. **c**, Population-averaged Fano factor of STN/GPe/GPi neurons for the short- and long-MT trials in the PRE and POST periods. Bin width, 100 ms. Error bars indicate SEM. * P < 0.05, ** P < 0.01, *** P < 0.001, two-tailed Wilcoxon signed rank test with Bonferroni correction (n = 44 STN, 79 GPe, and 78 GPi neurons).

To examine how STN suppression reduces cortical inputs to the GPe/GPi through the STN, neuronal responses to cortical stimulation in the STN and GPe/GPi were recorded (Supplementary Fig. 6). During the PRE period, the typical response of GPe/GPi neurons was triphasic, consisting of early excitation, inhibition, and late excitation phases, conveyed through the cortico-STN-GPe/GPi, cortico-striato-GPe/GPi, and cortico-striato-GPe-STN-GPe/GPi pathways, respectively^8,34^. During the POST period, early excitation was diminished, and inhibition enhanced (Supplementary Fig. 6a-c *middle* and *bottom*), suggesting that excitatory inputs from the STN to the GPe/GPi were reduced and that inhibitory inputs from the striatum were relatively enhanced. Baseline activity decreased significantly in the GPe but not GPi. Hence, STN suppression reduced the efficiency of information transmission from the cortex to the GPe/GPi via the STN as well as the baseline activity of the GPe.

**Fig. 6.**
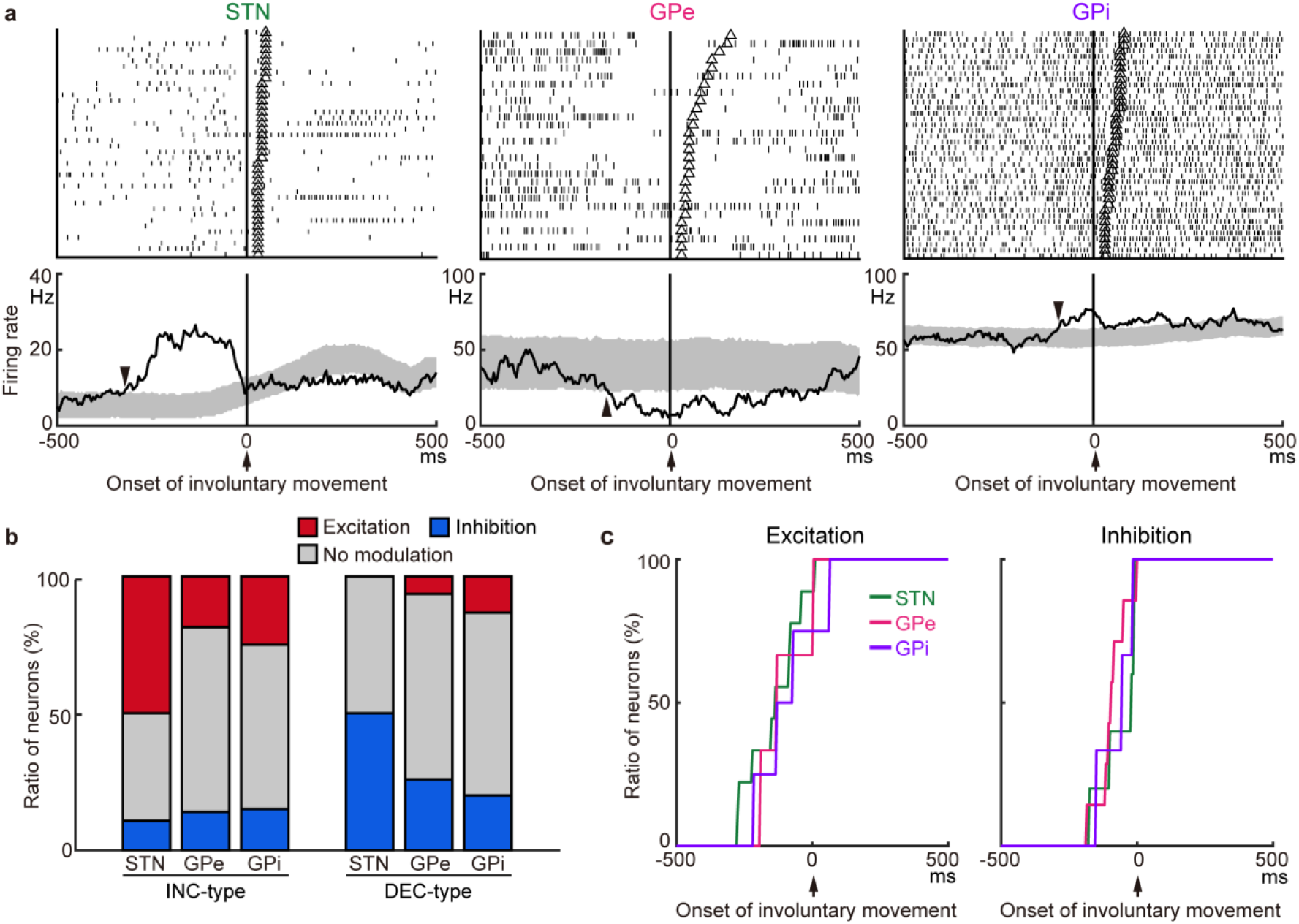
Neural activity in relation to involuntary movements after STN suppression. a, Typical examples of neural activity in the STN/GPe/GPi during involuntary movements. Spikes are aligned with the onset of involuntary movements, and trials are sorted by the duration of involuntary movements. Open triangles in the raster plots indicate the end of the involuntary movements. In the PETHs, a shuffling method was applied to estimate 95% confidence interval (shading), and the onset of activity modulation is indicated by arrowheads (−320, −160, and −90 ms for STN, GPe, and GPi neurons, respectively). Bin width, 5 ms; averaging window, 40 ms. **b**, Ratios of STN/GPe/GPi neurons exhibiting significant excitation or inhibition during involuntary movements among INC-type STN (n = 28 neurons), GPe (n = 43), and GPi (n = 20) neurons, and DEC-type STN (n = 10), GPe (n = 31), and GPi (n = 30) neurons. **c**, Cumulative histograms of the modulation onset timings for the STN/GPe/GPi neurons with excitatory (left) and inhibitory (right) modulations.

### GPe/GPi movement-related activity weakly affected by STN suppression

All GPe neurons with M1/SMA inputs exhibited movement-related activity during the PRE period and were thus classified as increasing (INC) type (46/79, 58%) or decreasing (DEC) type (33/79, 42%) based on the polarity of the largest movement-related modulation. GPe neurons in each type are exemplified in Supplementary Figure 7a, b. The activity of GPe neurons was summarized as heat maps (Fig. 3a) and population peri-event time histograms (PETHs) (Fig. 3b). In the POST period, the firing rate decreased in both types of GPe neurons, but the temporal structure of the movement-related modulation was preserved, i.e., INC- and DEC-type neurons showed increased and decreased activity, respectively, during movements. Statistical analyses revealed that the onset timing of movement-related modulation was similar for each neuron type between the PRE and POST periods (Fig. 3c3; INC type, from 33 ± 21 ms to 29 ± 19 ms, P = 0.9, n =46; DEC type, from 13 ± 27 ms to 34 ± 27 ms, P = 0.1, n = 33; Wilcoxon signed rank test; mean ± SEM). However, the following changes were observed in the POST period: 1) The baseline firing rate decreased (Fig. 3c1; INC, from 69.6 ± 4.0 Hz to 59.5 ± 4.1 Hz, P < 0.001; DEC, from 86.0 ± 5.2 Hz to 71.4 ± 5.3 Hz, P < 0.001); 2) The peak and trough amplitudes of the PETHs decreased (Fig. 3c2; peak amplitude in INC, from 66.7 ± 4.2 Hz to 57.5 ± 4.8 Hz, P < 0.05; trough amplitude in DEC, from 61.5 ± 4.3 Hz to 52.4 ± 4.7 Hz, P < 0.01); and 3) Movement-related modulation was prolonged (Fig. 3c4; INC, from 186 ± 25 ms to 312 ± 49 ms, P < 0.01; DEC, from 238 ± 34 ms to 318 ± 47 ms, P < 0.05).

Similarly, all GPi neurons with M1/SMA inputs exhibited movement-related activity in the PRE period and were classified as either INC type (32/78, 41%) or DEC type (46/78, 59%). GPi neurons in each type are exemplified in Supplementary Figure 7c, d. The activity of GPi neurons was also summarized as heat maps (Fig. 3d) and population PETHs (Fig. 3e). Movement-related activity was affected only weakly by STN suppression. Statistical analyses indicated that the following parameters did not change during the POST periods: 1) baseline firing rate (Fig. 3f1; INC, from 77.4 ± 5.3 Hz to 75.4 ± 6.1 Hz, P = 0.4, n = 32; DEC, from 88.4 ± 4.0 Hz to 87.4 ± 3.7 Hz, P = 0.4, n = 46); 2) PETH trough amplitude of DEC-type neurons (Fig. 3f2; from 62.6 ± 3.5 Hz to 62.6 ± 3.9 Hz, P = 0.5); 3) onset timing of movement-related modulations (Fig. 3f3; INC, from 0 ± 24 ms to −5 ± 23 ms, P = 0.5; DEC, from 27 ± 21 ms to −13 ± 19 ms, P = 0.3); and 4) duration of movement-related modulations in INC-type neurons (Fig. 3f4; from 241 ± 42 ms to 313 ± 59 ms, P = 0.3). The following parameters were exceptional: 1) PETH peak amplitude of INC-type neurons (Fig. 3f2; from 73.1 ± 6.1 Hz to 60.0 ± 6.4 Hz, P < 0.01); and 2) movement-related modulations in DEC-type neurons (Fig. 3f4; from 198 ± 21 ms to 235 ± 27 ms, P < 0.05).

The same analyses were performed for STN neurons (Supplementary Fig. 8). All STN neurons with M1/SMA inputs exhibited movement-related activity in the PRE period and were classified as INC type (32/44, 73%) or DEC type (12/44, 27%). Both types exhibited significantly reduced baseline firing rates, but movement-related activity was affected only weakly.

The above analyses with the PETH showed that the activity of GPe/GPi neurons was affected only weakly by STN suppression. Trial-averaging analyses such as PETHs may not be appropriate to explain trial-to-trial variability in task performance. Hence, the temporal structures of spike trains (e.g., bursts and pauses) were analyzed in detail.

### Spike train variability increased in STN and GPe/GPi neurons

Figure 4a shows examples of bursts and pauses in STN/GPe/GPi neurons. In the POST period, the pauses (blue lines) tended to become more frequent and of longer duration both at rest and during movements in all three examples, and burst activity (red lines) became more frequent in GPe neurons. Statistical analyses at rest and during movements revealed that the pause probability increased in the STN/GPe/GPi during the POST period (Fig. 4b; STN, from 18.5 ± 2.9% to 23.6 ± 3.4%, P < 0.05, n = 44; GPe, from 19.6 ± 1.7% to 27.4 ± 2.1%, P < 10^−7^, n = 79; GPi, from 7.7 ± 1.1% to 10.3 ± 1.3%, P < 10^−4^, n = 78; Wilcoxon signed rank test; mean ± SEM), whereas the burst probability increased only in GPe neurons (from 0.29 ± 0.05% to 0.61 ± 0.11%, P < 0.001). The coefficient of variation of inter-spike intervals (CV of ISIs) increased in the STN/GPe/GPi during the POST period (Fig. 4b; STN, from 1.03 ± 0.06 to 1.25 ± 0.06, P < 10^−4^; GPe, from 1.14 ± 0.05 to 1.50 ± 0.07, P < 10^−12^; GPi, from 0.89 ± 0.03 to 1.00 ± 0.04, P < 10^−6^). The sequential correlation (i.e., the correlation in spike count between two successive windows [window size, 20 ms]), which quantifies the temporal dependence of spikes^35^, increased in the STN/GPe/GPi during the POST period (Fig. 4b; STN, from 0.055 ± 0.031 to 0.114 ± 0.029, P < 10^−4^; GPe, from 0.298 ± 0.020 to 0.377 ± 0.022, P < 10^−9^, n = 79; GPi, from 0.169 ± 0.016 to 0.224 ± 0.015, P < 10^−6^, n = 78). The Fano factor, a measure of the variability in spike activity across trials, increased in the STN/GPe/GPi during the POST period (Fig. 4b; 100-ms bin size; STN, from 1.19 ± 0.10 to 1.52 ± 0.12, P < 10^−5^; GPe, from 1.63 ± 0.09 to 2.22 ± 0.13, P < 10^−11^; GPi, from 1.13 ± 0.07 to 1.31 ± 0.07, P < 10^−4^).

To investigate the neural mechanism that contributed to the Fano factor increase, its correlations with the firing rate and sequential correlation were examined (Fig. 4c). The changes in the Fano factor were negatively correlated with changes in the firing rate in the STN (Fig. 4c *upper*; *R*^*2*^ = 0.254) but not in the GPe/GPi (GPe, *R*^*2*^ = 0.141; GPi, *R*^*2*^ = 0.157). In contrast, the changes in the Fano factor were positively correlated with changes in the sequential correlation in all three nuclei (Fig. 4c *lower*; STN, *R*^*2*^ = 0.666; GPe, *R*^*2*^ = 0.578; GPi, *R*^*2*^ = 0.662). The same analysis was applied to other statistical measures (Fig. 4d). In addition to the sequential correlation, changes in the CV of ISIs were positively correlated with Fano factor changes in all three nuclei. Interestingly, changes in the pause probability were positively correlated with Fano factor changes in the GPe/GPi but not the STN. The pauses in the GPe/GPi became more frequent and of longer duration both at rest and during movements in the POST period (Fig. 4e). These results suggest that STN suppression induces sporadic pauses and interrupted spike trains, resulting in highly irregular, unstable neural activity.

### GPe/GPi spike variability is correlated with disturbance of reaching movements

Although STN suppression increased spike train variability (Fig. 4) in the GPe/GPi, it is not clear how these changes disturbed reaching movements. Detailed observations of spike trains and MTs (Fig. 5a) suggested that the firing rate of STN neurons tended to be lower in trials with long MTs in both the PRE and POST periods and that the Fano factor of GPe/GPi neurons tended to be higher in trials with long MTs. Thus, trials were grouped as short-MT trials (green, trials with MTs below the 25^th^ percentile) or long-MT trials (magenta, trials with MTs above the 75^th^). Population averaged PETHs revealed that the firing rate of the STN in the POST period from −300 to −200 ms from Movement onset was significantly lower in the long-MT trials, whereas no correlation between firing rate and MT was observed in the GPe/GPi (Fig. 5b). In contrast, population averaged Fano factors revealed that the Fano factor was higher in the long-MT trials during the POST period in the STN/GPe/GPi (Fig. 5c): at 0 ms from Movement onset in the STN, from 0 to 300 ms in the GPe, and from −100 to 100 ms in the GPi. Interestingly, the Fano factor tended to be higher in the long-MT trials during the PRE period as well, suggesting that STN suppression exaggerates spike train variability observed in the normal state.

### Involuntary movements correlated with phasic STN, GPe, and GPi activity

The relationship between phasic activity changes in STN/GPe/GPi neurons and involuntary movements was examined in the POST period (Fig. 6). Among neurons with a sufficient number of trials (>20) with involuntary movements, STN (22/38, 58%), GPe (24/74, 32%), and GPi (18/50, 36%) neurons exhibited significant firing rate modulations during the 200 ms preceding the onset of involuntary movements, as shown in Figure 6a (data shuffling method with α = 0.05, two-tailed; see Online Methods). In the STN, DEC-type neurons tended to exhibit inhibition during involuntary movements, and INC-type neurons tended to exhibit excitation (Fig. 6b; INC vs. DEC types, *χ*^2^ = 10.6, P < 0.005; chi-square test). A similar tendency was observed in the GPe (although only marginally significant: *χ*^2^ = 3.3, P = 0.07) but not the GPi (*χ*^2^ = 1.16, P = 0.3). Both the excitation and inhibition preceded the onset of involuntary movements (Fig. 6c): excitation (STN, −138 ± 100 ms; GPe, −105 ± 100 ms; GPi, −87 ± 118 ms; mean ± SD) and inhibition (STN, −63 ± 73 ms; GPe, −91 ± 57 ms; GPi, −73 ± 69 ms).

## Discussion

Simplified neural networks with excitatory and inhibitory neurons can generate irregular spike patterns without external noise, either through fluctuations in synaptic inputs around the spike threshold^36^ or through shifts in network states^37,38^. These models require interactions between excitatory and inhibitory neurons. The reciprocal connection between the STN and GPe is sufficient to generate irregular spiking patterns, which could be transferred to the GPi. One possible mechanism involves frequent shifts in STN-GPe network dynamics. With the reduced excitatory tone, the population activity can more easily shift from one network state to another. Such a frequent network transition would result in varying spike counts at the same task timings across trials, leading to an increase in the Fano factor. Neural mechanisms that modulate spike train variability via excitatory external inputs have been proposed; simulation studies showed that simple or correlated excitatory inputs to a neural network reduce spike variability^37,39^, which could explain the high spike variability in the GPe/GPi upon STN suppression. Although the exact neural mechanism is not clear, the STN-GPe reciprocal connection is suitable to control irregularities in spike trains.

Exaggerated pauses in the GPe can also contribute to development of spike variability (Fig. 4). The functional role of pauses in the GPe remains elusive, however. Many GPe neurons reportedly exhibit pauses in non-human primates^33,40^ and humans^41^. Although pauses were not correlated with any movements, relationships with alertness, task engagement, and motor learning have been reported^42–44^. Hence, the probability of pauses depends upon the animal’s state, while the timing of individual pauses does not have any physiologic significance. Loss of excitatory inputs from the STN to the GPe dramatically enhanced pauses (Fig. 4e), consistent with pharmacologic studies^8,34^. Assuming that the GPe affects the GPi either directly or indirectly, the random and irregular nature of pauses in the GPe is sufficient to impart variability to spike trains in the GPi.

Abnormal involuntary movements induced by STN suppression occurred in a sporadic manner irrespective of task timing or type (Fig. 2d). Compared with hemiballism induced by lesions or complete blockade of the STN^6–8^, DREADD-induced involuntary movements exhibited a smaller amplitude and affected only a specific body part, presumably because suppression of STN activity was mild (65-75%) and affected only the forelimb motor subregion.

After STN suppression, spike variability increased in STN/GPe/GPi neurons (Fig. 4), some of which exhibited activity modulation preceding the onset of involuntary movements (Fig. 6). These results suggest that involuntary movements are induced via the following neural mechanism: 1) In the normal state, excitatory inputs from the STN stabilize GPe/GPi activity; 2) Loss of excitatory inputs from the STN increases pauses and spike variability in the GPe/GPi; and 3) Coincident pauses/spikes across neurons occur with an increased probability, leading to sporadic involuntary movements. Neural activity in the STN/GPe/GPi exhibited similar modulation (i.e., increase or decrease) between in involuntary movements and during the task (Fig. 6b), and such modulation well preceded the onset of involuntary movements (Fig. 6c). These observations suggest that involuntary movements are conveyed through the same cortico-BG pathway that regulates normal voluntary movements.

The STN is believed to increase the baseline firing rate of the GPi, enhancing inhibition on the thalamus. The role of the STN in regulating movement stopping or switching was initially examined in DBS and imaging studies in humans^20–22^, and later supported by electrophysiologic recordings in rodents^45^. Further studies in non-human primates showed that the STN neurons involved in movement stopping or switching are restricted to the ventromedial STN^46,47^, the target of the dorsolateral prefrontal cortex or pre-supplementary motor area^48,49^. The present study focused primarily on the dorsolateral STN (Fig. 1b and Supplementary Fig. 1), the target of the M1 and SMA^29,48,50^. Together with the motor-related STN activity observed during the reaching task (Supplementary Fig. 8), the dorsolateral STN plays a role more closely related to the movement itself rather than movement stopping or switching.

Our across-trial analysis showed that long MT was associated with high spike variability at Movement onset after STN suppression in STN/GPe/GPi neurons, but no such trend was observed in terms of firing rate (Fig. 5). Interestingly, low STN firing rates preceded high spike variability in GPe/GPi neurons by 200-300 ms (Fig. 5b, c), supporting a role for the STN in maintaining stability in the BG circuitry. These results suggest a novel perspective on the STN function; Excitatory STN inputs stabilize spike timing in GPe/GPi neurons, reduce trial-to-trial variability during movements, and contribute to performance of rapid and stable movements. Loss of such excitatory inputs increases spike variability in GPe/GPi neurons and disturbs movements.

Loss of dopaminergic neurons in the substantia nigra pars compacta results in motor impairments in PD. The classical model predicts that diminished transmission via the *direct* pathway and enhanced transmission via the *indirect* pathway lead to increased GPi activity^5^; however, the baseline firing rate in the GPi does not necessarily increase^9,51,52^. Instead, the firing pattern changes dramatically in PD; spike trains of many GPe/GPi neurons exhibit correlated, oscillatory activity at the β frequency^53,54^. Although the origin of these pathologic oscillations remains controversial, the STN-GPe reciprocal loop is reportedly crucial^13,14^. Both STN-DBS and STN suppression may antagonize these GPe/GPi activity changes observed in PD. Actually, both DBS and lesions of the STN exhibit therapeutic effects on PD symptoms^15–17,55^. High-frequency DBS inhibits STN activity, presumably by stimulating GABAergic presynaptic terminals^56–58^, activates STN axons and induces a regularized and phasic-locked firing pattern in GPe/GPi neurons, resulting in a reduction in information transmission among the BG nuclei^58–61^. Chemogenetic suppression of the STN increased pause frequency and duration and the sequential correlation in the STN/GPe/GPi (Fig. 4), indicating that the discharge rate of a neuron depends more on its previous state than the input to the neuron. Therefore, STN-DBS and STN lesions including chemogenetic suppression would have the same effect: a reduction in the transmission of information across the BG nuclei, which could prevent the spread of pathologic oscillatory activity within and/or outside the BG and thus have beneficial effects on PD symptoms.

## Online Methods

### Experimental subjects

Three Japanese monkeys (*Macaca fuscata*; E, male, 7.9 kg, 6 years old at the time of surgical operation; K, female, 6.7 kg, 7 years old; U, female, 5.1 kg, 4 years old) were used in this study. During behavioral experiments, access to drinking water was restricted to maintain body weight at 90% of initial and then completely withheld for 24 h before experiments. The experimental protocols were approved by the Institutional Animal Care and Use Committee of the National Institutes of Natural Sciences. All experiments were conducted according to the guidelines of the National Institutes of Health *Guide for the Care and Use of Laboratory Animals*.

### Surgery

Each monkey underwent surgical operation under aseptic conditions to fix its head painlessly in a stereotaxic frame, as previously described^8,62^. Under general anesthesia with ketamine hydrochloride (5-8 mg/kg body weight, i.m.), xylazine hydrochloride (0.5-1 mg/kg, i.m.), and propofol (5-7 μg/ml of target blood concentration, i.v.), the scalp was incised, the skull was widely exposed, and bolts made of polyether ether ketone (PEEK) or titanium were screwed into the skull as anchors. The skull was covered with bone adhesive resin (Super-Bond C&B, Sun Medical) followed by acrylic resin (Unifast II, GC Co). Two PEEK pipes were mounted in parallel over the frontal and occipital areas for head fixation. An antibiotic was injected (i.m.) after surgery.

After one week of recovery time, bipolar stimulating electrodes were chronically implanted to the motor cortices^8,28,62^. Under general anesthesia with ketamine hydrochloride (4-5 mg/kg, i.m.), the skull over the forelimb regions of the M1 and SMA was removed. To access the STN vertically or the GPe/GPi obliquely, the skull on the trajectories was also removed. The forelimb regions of the M1 and SMA were physiologically identified^8,62^. Three pairs of bipolar stimulating electrodes (200-μm stainless steel wires; 2-2.5 mm inter-electrode distance) were then implanted in the distal and proximal forelimb regions of the M1 and the forelimb region of the SMA and fixed with acrylic resin: two pairs to distal and proximal forelimb regions of the M1, and one pair to the forelimb region of the SMA. Rectangular plastic chambers were fixed to the skull with acrylic resin to access the STN and GPe/GPi. In monkey E, the stimulating electrode in the SMA became ineffective during experimental sessions, and only the stimulating electrodes in the M1 were used.

### Preparation of AAV

The transfer plasmid, pAAV-CAG-hM4D-2a-GFP (Fig. 1a), was prepared from pAAV-CAG::FLEX-rev::hM4D-2a-GFP, a gift from Scott Sternson (Addgene plasmid #52536)^63^. The DNA fragment encoding hM4D-2a-GFP was separated at two *Eco*RI sites and inserted into the original plasmid in the inverted orientation, followed by the excision of the FLEX (loxP and lox2272) sequence at *Xba*I and *Spe*I sites.

The AAV vector was prepared as previously described^64^. Briefly, the plasmid vector was packaged with AAV-DJ capsid using the AAV Helper Free Expression System (Cell Biolabs); the packaging plasmids (pAAV-DJ and pHelper) as well as the transfer plasmid were transfected into HEK293T cells, which were harvested 72 h later and lysed by repeated freezing and thawing. The crude cell extract containing AAV particles was purified by ultracentrifugation with cesium chloride and concentrated by ultrafiltration using an Amicon 10K MWCO filter (Merck Millipore). The copy number of the viral genome (vg) was 6.5-9.5×10^12^ vg/ml, as determined using TaqMan Universal Master Mix II (Applied Biosystems).

### Mapping the STN

Extracellular unit activity was recorded with glass-coated tungsten electrodes (1 MΩ, Alpha Omega) or handmade Elgiloy-alloy microelectrodes (0.5-1.5 MΩ at 1 kHz). A microelectrode was inserted vertically into the STN. Signals from the electrode were amplified, digitized at 44 kHz, digitally filtered between 0.5 and 9 kHz, and stored on a computer using a multi-channel recording system (AlphaLab SnR, Alpha Omega). A custom MATLAB script was used to manually isolate single-unit activity of the STN neurons. The STN was identified based on (1) mid-frequency (40 Hz) firings, and (2) biphasic excitation to cortical stimulation (Fig. 1a; 0.3-ms duration; single pulse; intensity, 0.2 to 0.7 mA; inter-stimulus interval, 1.4 s) examined by constructing peri-stimulus time histograms (PSTHs) with 1-ms bins^8,28,62^.

### Injection of AAV

To precisely target the motor subregion of the STN, neural activity was recorded using a micropipette with a wire electrode when exploring the AAV injection sites. A glass micropipette was made from a borosilicate glass capillary (inner diameter, i.d., 1.8 mm; outer diameter, o.d., 3 mm.; G-3, Narishige) using a puller (PE-2, Narishige) and a beveler (EG-3, Narishige) and connected to a 25-μl Hamilton microsyringe (Hamilton Company) by a joint Teflon tube (JT-10, Eicom). A tungsten wire (30-μm core diameter with Teflon insulation; California Fine Wire Co.) was inserted into the micropipette to record neuronal activity. The glass micropipette, tubing, and Hamilton microsyringe were filled with mineral oil (MOLH-100, Kitazato Co.). The syringe was mechanically controlled by a syringe pump (IMS-20, Narishige). Viral vector solution was loaded from the micropipette. The glass micropipette was inserted vertically into the motor subregion of the STN through a small incision in the dura mater based on the STN mapping. After confirming the motor subregion of the STN by responses to cortical stimulation, 1 μl of the AAV solution was slowly infused at a rate of 0.05 nl/min. The micropipette was left in place for an additional 5 min and then slowly moved. To cover the motor subregion of the STN, multiple injections were performed (1 μl per site, 1-2 sites per track, 1-2 tracks per day for 2-4 days); in total, 4, 15, and 21 μl were injected to the STN of monkeys E, K, and U, respectively.

### Reaching task

Each monkey was trained to sit on the monkey chair and perform a custom reach-and-pull task, in which the monkey initiated the movements immediately after a ‘Go’ cue presentation (externally triggered reaching trials, or ET trials) or without an apparent Go cue (self-initiated reaching trials, or SI trials) (Fig. 2a). Task setup consisted of a home lever (2.5 cm in length, located 20 cm away and 25 cm below eye position), a full color LED (located at 25 cm away and 5 cm below), and a front lever (located at 22, 22, and 20 cm away for monkeys E, K, and U, respectively, and 7 cm below). First, the monkey sat on the monkey chair with its head fixed and pulled the home lever toward its body. After a random delay of 0.8-2.5 s, the LED turned on (Task cue) with a color instructing the trial type (blue, ET trial; red, SI trial). In ET trials, the LED color changed from blue to green (Go cue) in 1-2 s; the monkey was required to release its hand from the home lever (Movement onset) within 0.5 s and pull the front lever (Lever pull) within 3 s. In SI trials, the monkey was required to wait for 1.5 s but no more than 5 s, release its hand from the home lever (Movement onset), and pull the front lever (Lever pull) within 3 s. The trials were considered successful only if both the home lever release and the front lever pull were performed within the correct time windows. With a delay of 0.5 s after the front lever pull, the LED turned off; in a successful trial, 0.2 ml of juice was delivered as a reward. The two types of trials were randomly presented with an equal probability of 45%. The remaining 10% of trials were the same as SI trials except that the red LED turned green at 1.5 s to instruct the monkey as to the correct timing of the movement initiation (Instruction trials). The positions of the home and front levers were monitored using magnets attached to the levers and Hall-effect sensors fixed on the lever housings. Analog outputs from the sensors were recorded at 2,750 Hz, down-sampled to 100 Hz, and converted to the lever positions.

RT was defined as the time from Go cue to Movement onset in ET trials and the time from Task cue to Movement onset in SI trials. MT was defined as the time from Movement onset to Lever pull. The behavioral task was controlled and logged using a custom script written in LabVIEW (LabVIEW 2013, National Instruments).

Stable performance was achieved in all monkeys after training for >3 months; the success rates were 95.1 ± 2.9, 88.7 ± 10.1, and 89.2 ± 10.5% in ET trials and 82.1 ± 10.1, 78.8 ± 7.4, and 91.6 ± 7.7% in SI trials for monkeys E, K and U, respectively (mean ± SD; n = 21, 28, and 32 for monkeys E, K, and U, respectively). To avoid possible effects of DREADD ligands from the previous experiment, task sessions were performed every other day.

### Administration of CNO or DCZ

CNO (HY-17366, MedChem Express) and DCZ (HY-42110, MedChem Express) were dissolved in dimethyl sulfoxide (DMSO) and then diluted with 0.9% saline to a final concentration of 1 mg/ml in 5% DMSO solution. DCZ is a newly developed ligand with high *in vivo* stability and high blood-brain-barrier permeability^32^, without the potential off-target effects associated with CNO^65^. Aliquots were frozen at −20°C for <2 weeks until used. The amount of DREADD ligand was determined in a pilot study to induce abnormal movements and used throughout the experiments: CNO, 1.0 mg/kg body weight (i.v.) and DCZ, 0.1 mg/kg body weight (i.m.). The PRE period was defined as −15 to 0 min relative to administration of DREADD ligand. The POST period was defined as 20 to 45 min for CNO administration and 10 to 45 min for DCZ administration based on observations of STN activity (Fig. 1c), reflecting more rapid onset of DCZ than CNO. As a control, the same volume of vehicle (5% DMSO in 0.9% saline) was administered (i.v. in monkey E; i.m. in monkeys K and U).

### Behavioral observations

To reconstruct 3D trajectories for arm joints, RGB (x-y) and depth (z) images of the upper limb of the monkey during task performance were captured at 30 Hz using a depth camera (RealSense™ D435, Intel) and stored on a computer. To detect the positions of the arm joints, RGB (x-y) images were processed using DeepLabCut, as described previously^66,67^, using a computer (Ubuntu 18.04, Intel Core i7-8750H and GeForce GTX 1050Ti). Training data were prepared by manually labeling the shoulder, elbow, wrist, and hand of 160 images from 4 task sessions, and the neural network was trained for 200,000 iterations. After determining the positions of the joints using a RGB (x-y) image, the distance from the camera to each joint was calculated as the average depth close to the corresponding position (≤10 pixels) in the depth image. Lastly, time series of 3D trajectories were constructed and digitally low-pass filtered (4th-order Butterworth, 7.5 Hz) before analysis. Trajectory deviation was defined as the total deviation from the mean trajectory; tortuosity was defined as the trajectory length divided by the end-point distance; and travel distance was defined as the total trajectory length. Speed of the joints was calculated from the displacement between two consecutive frames (1/30 s).

### Recording of STN, GPe, and GPi activity

The motor subregion in the GPe/GPi receiving input from the M1 and SMA was roughly mapped with extracellular unit recordings using the similar electrode for the STN mapping. The electrode was obliquely (40° from vertical in the front plane) inserted, and spontaneous firing activity and cortically evoked responses were recorded. The GPe/GPi were identified based on 1) high-frequency (>60 Hz) firings, and 2) an excitation-inhibition-excitation triphasic response to cortical stimulation (the same parameters as those used for STN mapping^8,28,62^. The GPe and GPi were distinguished by 1) the GPe-GPi boundary with low-frequency firings, presumably the medial medullary lamina; and 2) firing patterns: pauses observed in HFD-P GPe neurons but rarely seen in HFD GPi neurons^33,40^.

Extracellular unit activity during task performance was recorded using 16-channel linear electrodes (0.8-2.0 MΩ at 1 kHz; inter-electrode distance, 100 or 150 µm; Plexon). The multichannel electrode was inserted through a stainless guide tube (i.d., 0.45 mm; o.d., 0.57 mm) vertically into the STN or obliquely (40° from vertical) into the GPe/GPi. Signals from the electrodes were amplified, digitized at 44 kHz, digitally filtered between 0.5 and 9 kHz, and stored on a computer using a multi-channel recording system (AlphaLab SnR, Alpha Omega). In a total of 78 recording sessions, 1 to 5 well-isolated units (2.54 ± 1.11; mean ± SD) were simultaneously recorded. A custom MATLAB script was used to manually isolate single-unit activity of STN/GPe/GPi neurons. Cortically evoked responses (the same parameters as those used for STN/GPe/GPi mapping) of STN/GPe/GPi neurons were examined by constructing PSTHs with 1-ms bins, and neurons with cortical inputs were analyzed. Cortical stimulation was performed every 5 or 10 min during the reaching task to examine the effect of the DREADD ligand (Supplementary Fig. 6). Electrophysiologic data obtained during cortical stimulation were excluded in analyses of baseline and movement-related activity.

### EMG recording

EMGs of the biceps and triceps brachii muscles were obtained using surface electrodes (NE-05, Nihon Kohden) for monkey K or chronically implanted stainless wire electrodes (7-stranded 25.4-μm diameter wire with Teflon coating; A-M Systems, Sequim, WA, USA) for monkey U. Signals from the electrodes were amplified (5000×) and bandpass-filtered (150-3000 Hz) using an amplifier (MEG-5200, Nihon Kohden), digitized at 20 kHz (PCIe-6321, National Instruments), and stored on a computer. The root mean square (RMS) of each EMG was calculated with a 100-ms moving window. The RMS of the EMG was aligned with task events, and the mean and SEM were computed (Supplementary Fig. 4).

### Histology

Monkeys E and K were sacrificed 58 and 89 weeks after AAV injection to examine viral transduction and confirm the sites of electrophysiologic recordings. Monkey U is still alive and used for further experiments. At the end of the experiments, electrolytic lesions were made with cathodal constant current (20 μA for 30 s) at the putative boundaries of the GPe/GPi. With an overdose of sodium pentobarbital (50 mg/kg, i.v.), the monkeys were perfused transcardially with 0.1 M phosphate buffer (PB) containing 4% formaldehyde. The brains were removed, kept in 0.1 M PB containing 30% sucrose at 4°C for cryoprotection, and serially cut with a freezing, sliding microtome (HM440E, Microm Co.) to obtain 50-μm-thick frontal brain sections.

For double immunofluorescence staining, free-floating sections containing the STN were incubated with rabbit anti-GFP (1:1000; A11122, Invitrogen) and mouse anti-NeuN (1:1000; MAB377, Millipore) primary antibodies overnight at 4°C. The sections were then rinsed and incubated with Alexa Fluor 488–conjugated goat anti-rabbit (1:500; A11043, Invitrogen) and Alexa Fluor 594–conjugated goat anti-mouse (1:500; A11032, Invitrogen) antibodies for 1 h at room temperature. Similarly, free-floating sections containing the GPe/GPi were stained with mouse anti-NeuN primary antibody and Alexa Fluor 594–conjugated anti-mouse secondary antibody. The sections were mounted on a gelatin-coated slide glass with FluorSave reagent (Calbiochem). Fluorescence images were taken on an inverted microscope (BZ-X700, Keyence) with a 10× objective.

### Data analysis

All data were analyzed using custom scripts written in MATLAB (MATLAB R2019b, MathWorks). Only successful trials were analyzed, except for the reaction and movement times shown in Figure 2b. Single-unit recordings from the STN/GPe/GPi were qualitatively similar in the three monkeys; thus, the datasets from all monkeys were combined for the electrophysiologic analysis.

### Neuronal responses to cortical stimulation

To examine the effect of STN suppression, cortically evoked responses in STN/GPe/GPi neurons were analyzed (Supplementary Fig. 6). First, PSTHs with 1-ms bins were constructed for each neuron; the PSTHs were smoothed with a Gaussian distribution (σ = 1.6 ms) and transformed to z-scores with activity from the 100 ms preceding cortical stimulation. For GPe/GPi neurons, early excitation, inhibition, and late excitation were defined as 3 to 200 ms after stimulation, whereas only early and late excitations were defined for STN neurons. Two consecutive bins with z > 1.65 or z < −1.65 were considered the onset of excitation and onset of inhibition, respectively. The latency of the response was defined as the time from stimulation to onset, and duration was defined as the period from onset until the first bin, followed by two consecutive bins within the threshold (|z| ≤ 1.65). The amplitude was defined as the sum of the |z| scores during each response. Early and late excitation was defined as excitation with latencies of <20 ms and ≥20 ms, respectively.

### Movement-related neuronal activity

Raster plots were constructed by aligning spikes at each task event, usually Movement onset (Supplementary Fig. 7), and displayed chronologically before and after DREADD ligand administration. PETHs were constructed by averaging firing rate across trials in 10-ms bins in the PRE period (during the 15 min before DREADD ligand administration) and POST period (20-45 min after CNO administration or 10-45 min after DCZ administration). The spike activity during the 500 ms before the Task cue presentation was used to calculate the baseline firing rate.

Modulation onset was defined as the time first exceeding a threshold of the mean ± 1.96 SD of the baseline firing rate, and modulation duration was defined as the time from onset until the last bin exceeding the threshold. The amplitudes of peaks and troughs were defined as the difference in peak and trough firing rate from the baseline firing rate, respectively. If more than one peak and/or trough was detected, the modulation having the largest area exceeding the threshold was used for the statistical analysis and classification of response type (i.e., INC or DEC type). Heat maps and population PETHs were constructed by calculating z-scores of PETHs using the firing activity during −500 to −300 ms relative to Movement onset in the PRE period as the baseline.

### Firing patterns

Bursts and pauses were detected using the Robust Gaussian Surprise method^68^. Briefly, the ISIs of a spike train were log-transformed to give log(ISI)s. First, the central distribution of log(ISI)s was calculated by excluding outliers, that is, bursts and pauses. The E-center was defined as the midpoint of the 5th and 95th percentiles of the log(ISI)s. The central set was defined as the log(ISI)s that fell within E-center ± 1.64 × MAD, where MAD is the median absolute deviation of log(ISI)s. The C1-center was defined as the median of the central set. The Central Location (μ) was the median of the log(ISI)s that fell within C1-center ± 1.64 × MAD. Then, normalized log(ISI)s (NLISIs) were defined as: NLISI_i_ = log(ISI_i_) – μ. The distribution of NLISIs was assumed to be Gaussian, with σ = 1.48 × MAD, and the P-value for each ISI was computed. ISIs below or above a statistical significance level (P < 10^−5^) were defined as bursts and pauses, respectively. Bursts and pauses were extended to adjacent ISIs if their inclusion lowered the P-value.

To analyze the spike train variability in Figure 4, the following parameters were calculated from −1 to 1 s relative to Movement onset in both the ET and SI trials: firing rate, burst and pause probabilities (probabilities of a neuron being in bursts and pauses, respectively), pause rate (frequency of pause occurrence), pause duration (average durations of all pauses), CV of ISIs (standard deviation divided by the mean of the ISIs), sequential correlation (correlation of spike count between successive time windows in each trial), and the Fano factor (variance divided by the mean number of spikes in a 100-ms window).

### Involuntary movements

Involuntary movements of the upper limb were detected as the task-irrelevant movements of the home lever from −0.5 to 1 s relative to Task cue. In the POST period, the home lever position exceeding the mean ± 3SD calculated from the same duration in the PRE period was considered to be caused by involuntary movements. The onset and end of involuntary movements were defined as the first and last points exceeding the mean ± 3SD, respectively. The home lever position from −0.5 to 1.0 s from Task cue was used to calculate the occurrence probability (Fig. 2d, e), and that from 0.1 to 1.5 s before Movement onset was used to analyze neural activity (Fig. 6).

To analyze spike activity during the involuntary movements shown in Figure 6, PETHs during ±500 ms from the onset of involuntary movements were constructed with 5-ms bins and 40-ms averaging windows. Involuntary movements were observed at various task timings (Fig. 2d); the movement-related activity around Movement onset would affect the PETH. To exclude the effect of movement-related activity, shuffled PETHs were constructed from trials without any involuntary movements and compared with the original PETH. To construct a shuffled PETH, each trial in the original PETH was replaced with a 1-s segment of spike train without involuntary movements at the same timing relative to Movement onset. This process was repeated 1000 times to obtain 1000 shuffled PETHs. At each bin, the spike counts of the 1000 shuffled PETHs were sorted from the lowest to the highest. The confidence interval was defined as the spike count from the 25^th^ to the 976^th^ at each bin, corresponding to a significance level of 0.05. Neural activity was judged to be modulated if the total spike count during the 200 ms preceding the onset of involuntary movements was below the 25^th^ (inhibition) or above the 976^th^ (excitation) of the corresponding spike counts in the shuffled PETHs. Onset of activity modulation was defined as the beginning of ≥10 bins (≥50 ms) exceeding the confidence interval during ±500 ms from the onset of involuntary movements.

### Statistics

The success rate, RT, and MT, as well as the occurrence probability of involuntary movements, were calculated in each session, and the statistical significance was computed using the two-tailed Wilcoxon signed rank test between the PRE and POST periods (Fig. 2b, e). In the trajectory analysis, trials from 2-3 sessions were combined; trajectory deviation, tortuosity, and maximum speed were calculated in each trial and compared between the PRE and POST periods using the two-tailed Mann-Whitney *U* test (Fig. 2c).

The significance of decreases in STN activity was computed using the one-tailed Wilcoxon signed rank test with Bonferroni correction (Fig. 1c). To determine the significance of changes in the PETHs (Fig. 3c, f and Supplementary Fig. 8e), firing patterns (Fig. 4b, e), and neuronal responses to cortical stimulation (Supplementary Fig. 6c), the parameters of individual neurons were compared between the PRE and POST periods using the two-tailed Wilcoxon signed rank test. The statistical significance of differences in spike properties between the long- and short-MT trials was computed using the two-tailed Wilcoxon singed rank test with Bonferroni correction (Fig. 5).

## Data availability

The data that support the findings of this study are available from the corresponding author upon reasonable request.

## Code availability

The code to generate the results and the figures of this study are available from the corresponding authors upon reasonable request.

## Acknowledgments

We thank S. Sato, H. Isogai, N. Suzuki, K. Awamura, K. Miyamoto, T. Sugiyama, and R. Kageyama for technical support; and Y. Yamagata for her critical reading of the manuscript. This work was supported by MEXT KAKENHI (“Non-linear Neurooscillology”, 15H05873 to A.N.), JSPS KAKENHI (19KK0193 to A.N., 16K07014 to S.C., JP18K15340 to T.H.), JST CREST (JPMJCR1853 to S.C.), AMED (JP20dm0307005 and JP20dm0207050 to A.N.), and Takeda Science Foundation (to T.H.) grants. Japanese monkeys used in the present study were obtained through the National Bio-Resource Project (NBRP) “Japanese Monkeys” of AMED.

## Author Contributions

T.H., S.C., and A.N. designed the study; K.K. generated the viral vector; T.H. and S.C. performed the experiments; T.H. analyzed the data; T.H., S.C., and A.N. wrote the manuscript.

