## Supplementary information for "Subthalamic nucleus stabilizes movements by reducing neural spike variability in monkey basal ganglia: chemogenetic study"

**Supplementary Video 1 | Involuntary movements induced by STN suppression**

Abnormal involuntary movements observed in monkey E at 70 min after CNO (1.0 mg/kg, i.v.) administration.

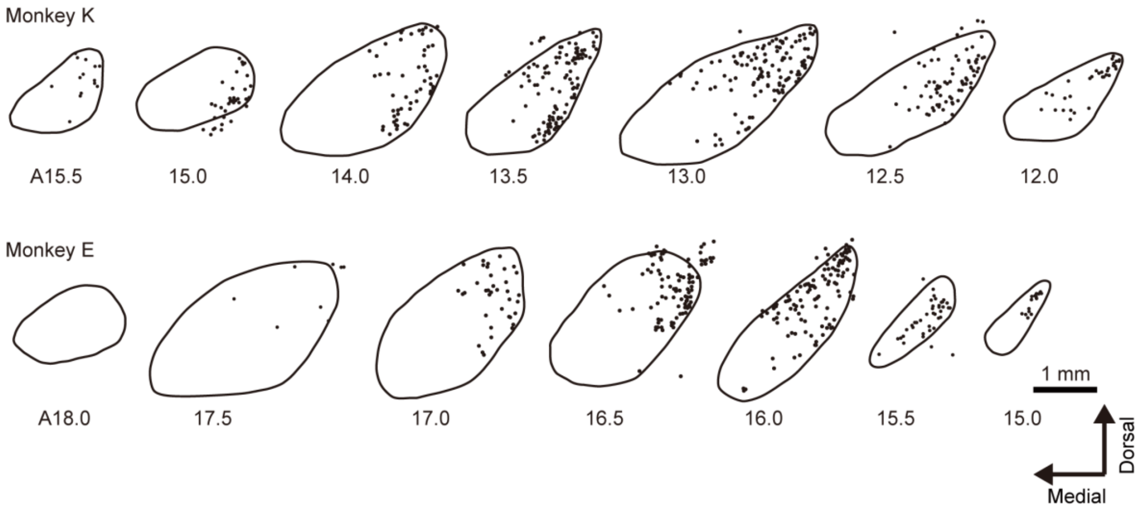


**Supplementary Fig. 1 | Distribution of transduced neurons in the STN.** Histologic examination of the STN of monkeys E and K. Cells labeled with a neuronal marker, NeuN, and anti-GFP antibody were considered transduced with AAV and indicated by black dots. Scale bar, 1 mm.

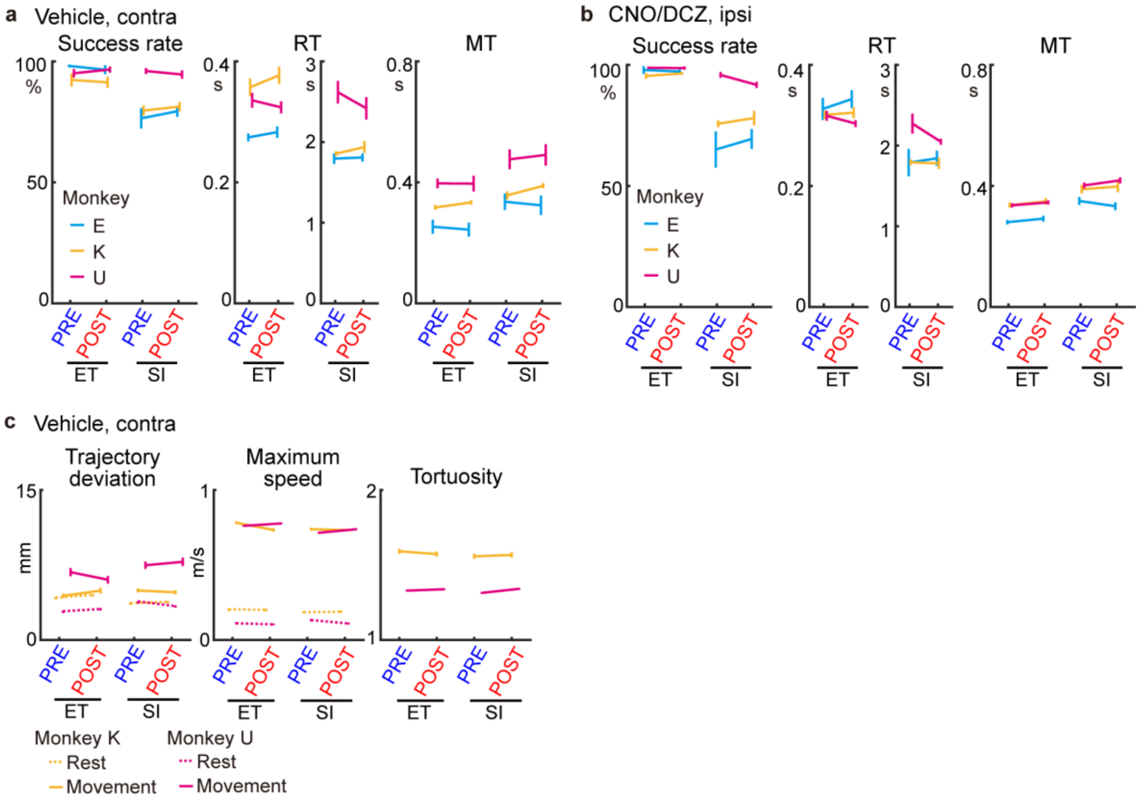


**Supplementary Fig. 2 | Control experiments for DREADD ligand administration. a**, **b**, Success rate, RT, and MT in control experiments, corresponding to Figure 2b. **a**, Performance of the task with vehicle administration. The hand contralateral ('contra') to the AAV injection side was used. Error bars indicate SEM. Two-tailed Wilcoxon signed rank test (n = 8, 12, and 10 sessions for monkeys E, K, and U, respectively). **b**, Performance of the task using the ipsilateral hand ('ipsi') with DREADD ligand administration. Error bars indicate SEM. Two-tailed Wilcoxon signed rank test (n = 8, 9, and 9 sessions). **c**, Analyses of wrist trajectories for monkeys K and U with vehicle administration, corresponding to Figure 2c. Error bars indicate SEM. Two-tailed Mann-Whitney *U* test (monkey K, 74 ET and 70 SI trials; monkey U, 45 ET and 54 SI trials).

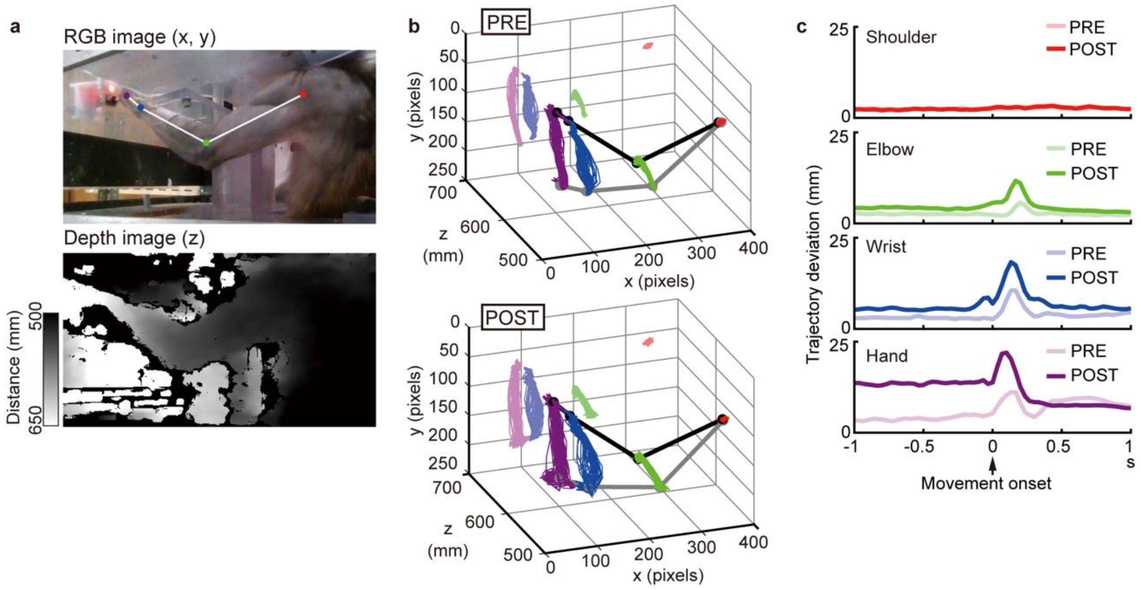


**Supplementary Fig. 3 | Trajectory analysis of arm joints during the task. a**, Example of RGB (x-y) and depth (z) images of monkey K captured using a depth camera. The RGB image was processed using DeepLabCut to detect the shoulder (red circle), elbow (green), wrist (blue), and hand (purple). **b**, Example 3D trajectories of the shoulder (red), elbow (green), wrist (blue), and hand (purple) from −1 to 1 s relative to Movement onset in ET trials of monkey K. Gray and black lines in the 3D plot indicate the mean positions at −1 and 1 s, respectively. Trajectories were projected on the x-y plane using pale colors. **c**, Trajectory deviations, the differences from the mean trajectories, in the PRE and POST periods.

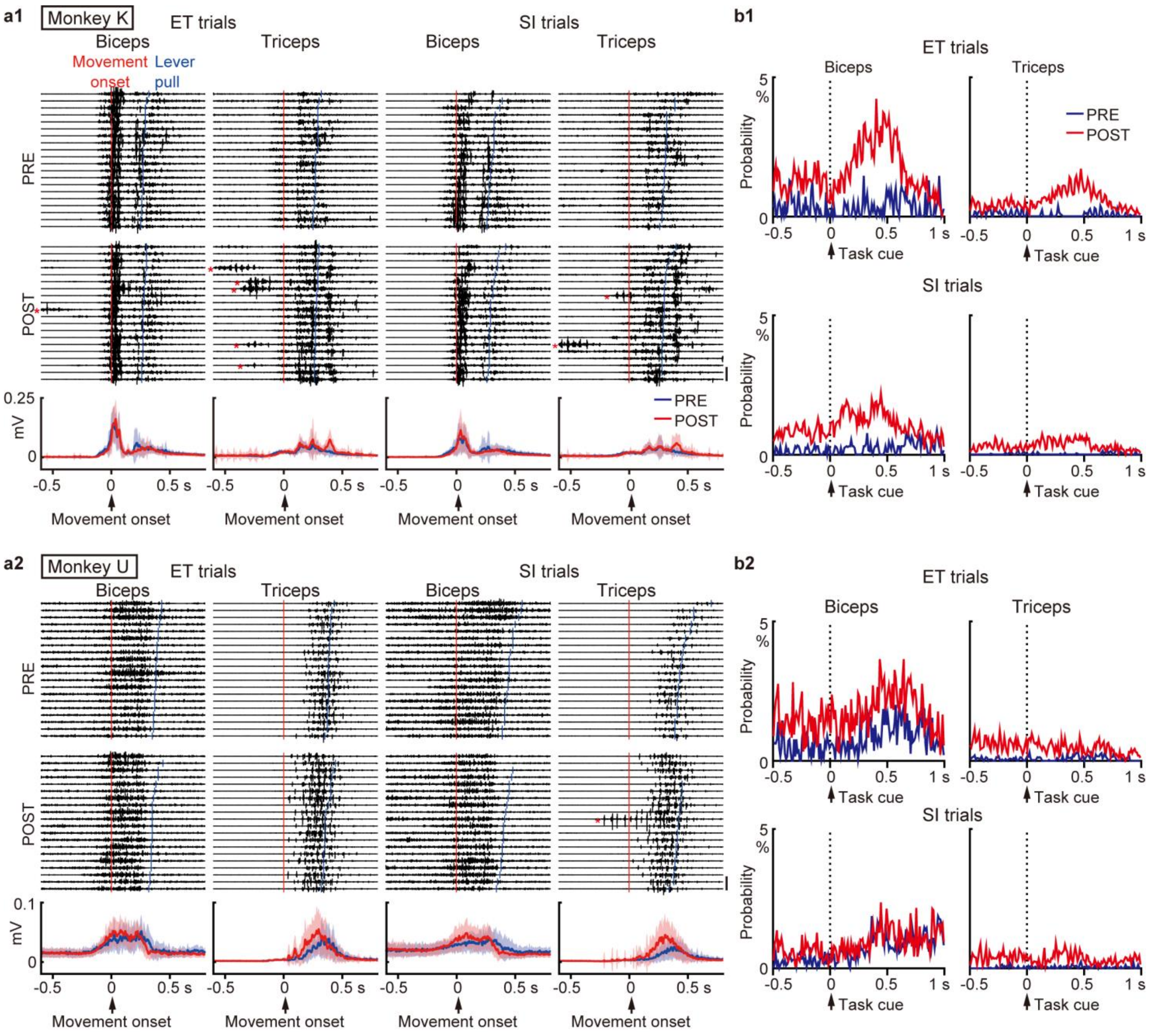


**Supplementary Fig. 4 | EMGs of the contralateral forelimbs of monkeys K and U during the task. a**, Raw EMGs and root mean square (RMS) of EMGs in a single session for monkey K (**a1**) and monkey U (**a2**), aligned with Movement onset in the PRE period (−15 to 0 min) and POST period (10 to 45 min). Examples of 20 EMG traces are sorted by MT (top two rows). Red and blue vertical lines indicate the timings of Movement onset and Lever pull, respectively. The RMS of EMGs was averaged in each period in 1-ms bins (mean ± SD; bottom). Red asterisks indicate task-irrelevant muscle activity, presumably corresponding to involuntary movements. Scale bar, 1 mV. Shading indicates SEM. **b**, Occurrence of abnormal EMG events before and after Task cue for monkey K (**b1**) and monkey U (**b2**). An abnormal EMG event was defined as an RMS increase above the mean + 3SD of the baseline activity (during the 500 ms before Task cue). Bin width, 10 ms. To calculate the occurrence probability, EMG recordings from 3 and 4 days were combined for monkeys K and U, respectively.

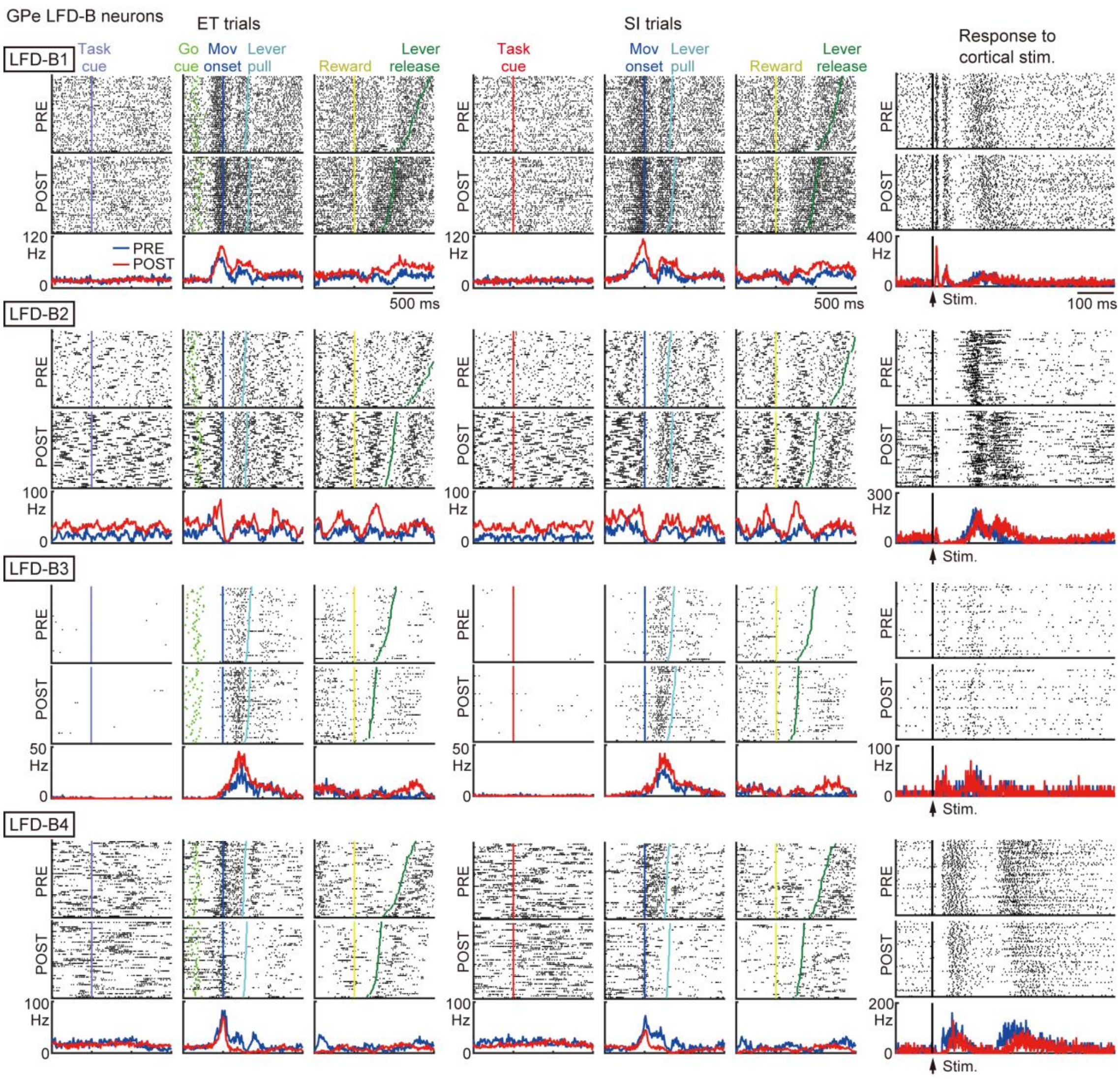


**Supplementary Fig. 5 | Raster plots, PETHs, and PSTHs to cortical stimulation of LFD-B neurons in the GPe.** Raster plots and PETHs of all LFD-B neurons in the GPe of monkey K. Spikes are aligned with Task cue, Movement onset (Mov onset), and Reward delivery in the ET and SI trials. Raster plots are sorted by the time to the next task event. Bin width, 10 ms. Four LFD-B neurons (1-4) responded to cortical stimulation (triphasic, inhibition, or excitation) and exhibited task-related activity. After STN suppression, task-related activity was generally enhanced (LFD-B1-3), and a burst firing pattern emerged. Further analysis of LFD-B neurons was not performed because of their small number. LFD-B and HFD-P neurons are considered to correspond to arkypallidal and prototypic neurons in the GPe reported in rodents[3].

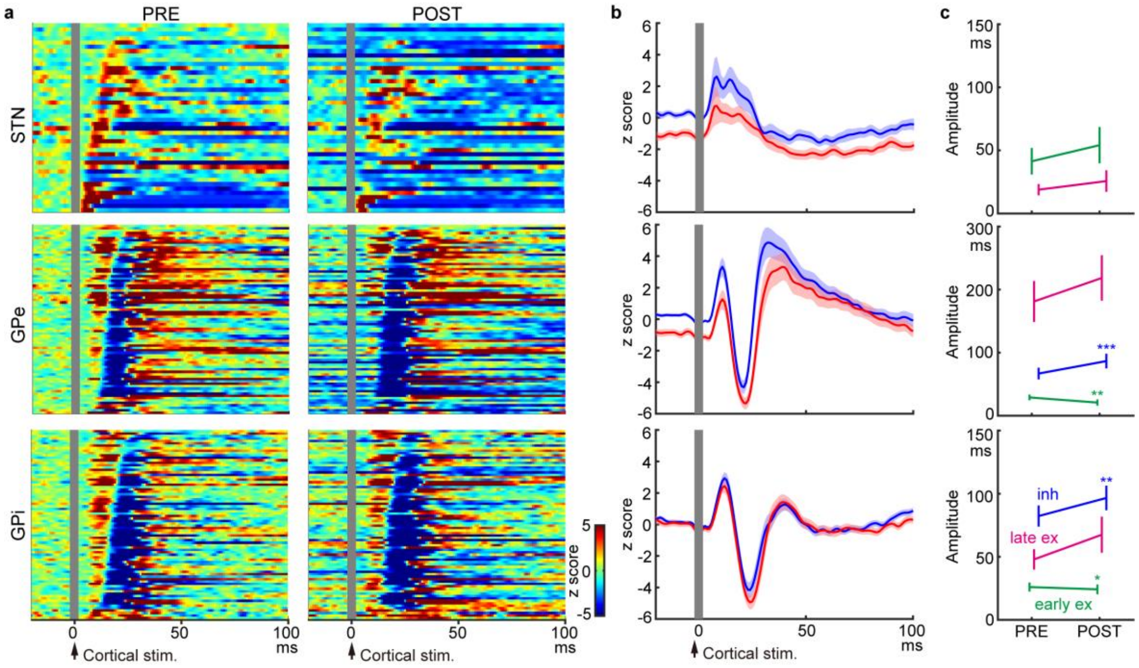


**Supplementary Fig. 6 | Cortically evoked responses of STN, GPe, and GPi neurons in the PRE and POST periods. a**, Heat maps of PSTHs of all STN (n = 44), GPe (n = 79), and GPi (n = 78) neurons in the PRE and POST periods. Each PSTH (width, 1 ms) was constructed from 100 repetitions of cortical stimulation, smoothed with a Gaussian filter (σ = 1.6 ms), and converted to z-scores using the 100 ms preceding stimulation in the PRE period. Neurons were sorted by the latency of earliest response in the PRE period for the STN and latency of inhibition for the GPe/GPi. Stimulation artefacts were covered by gray vertical bars at |t| < 2 ms. **b**, Population-averaged PSTHs of all STN/GPe/GPi neurons. Shading indicates SEM. **c**, Change in amplitude of early excitation (early ex) and late excitation (late ex) in STN neurons and early ex, inhibition (inh), and late ex in GPe/GPi neurons. Amplitude was defined as the area of the z-scored PSTH above z = 0 (early ex and late ex) or below z = 0 (inh). The baseline firing rate during the 100 ms preceding stimulation significantly decreased in GPe (from 77.0 ± 24.7 Hz to 65.4 ± 26.5 Hz, $P < 10^{-6}$; mean ± SD) but not GPi (from 84.9 ± 28.9 Hz to 82.5 ± 28.9 Hz, P = 0.3) neurons. Error bars indicate SEM. * P < 0.05, ** P < 0.01, *** P < 0.001, two-tailed Wilcoxon signed rank test.

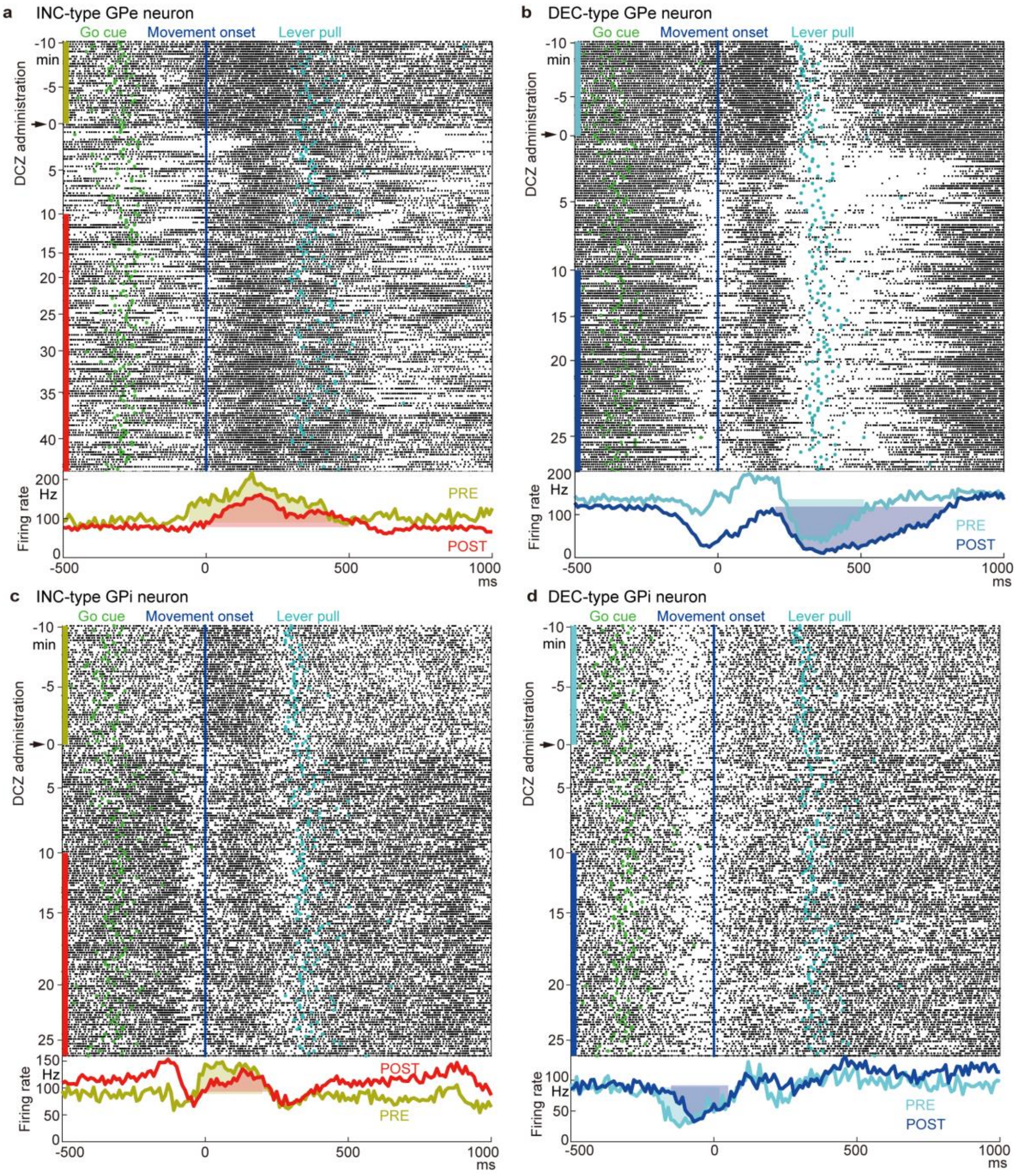


**Supplementary Fig. 7 | Examples of movement-related activity of GPe/GPi neurons. a**, Example of a GPe neuron exhibiting an activity increase (INC) during movements. Raster plot aligned at Movement onset is displayed chronologically by elapsed time during the experiment (top); vertical colored bars on the left indicate the PRE and POST periods. The timings of Go cue and Lever pull are also indicated. PETHs in the PRE and POST periods are plotted with different colors (bottom; 10-ms bins), with shading indicating movement-related modulations. **b**, Another GPe neuron demonstrating an activity decrease (DEC). **c**, **d**, Same as (**a**, **b**) but for GPi neurons.

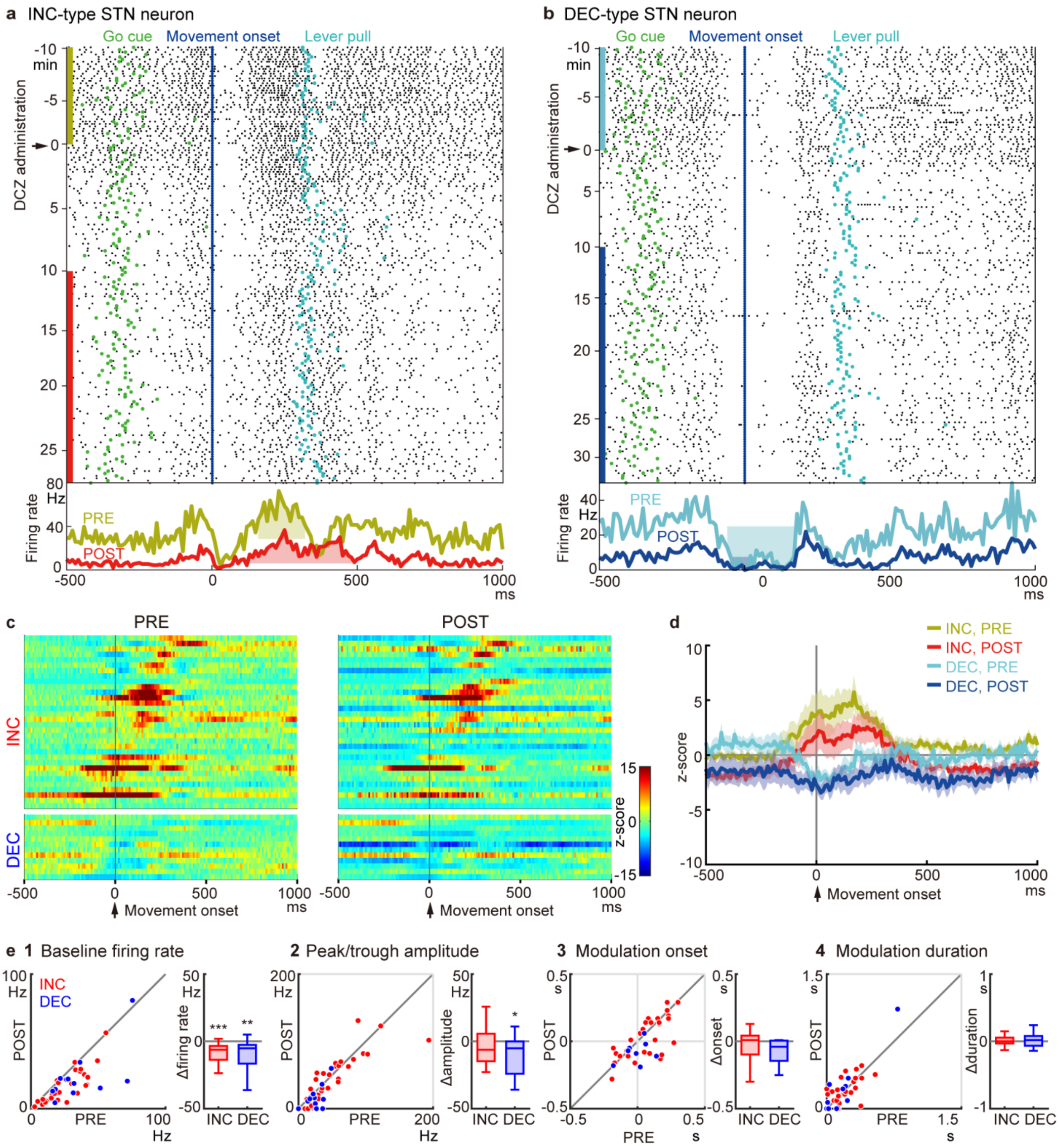


**Supplementary Fig. 8 | Changes in movement-related activity of STN neurons after STN suppression. a**, Example of a STN neuron exhibiting an activity increase (INC) during movements. **b**, Another STN neuron exhibiting an activity decrease (DEC). **c**, Heat maps for STN neurons classified as INC or DEC type. **d**, Population-averaged PETHs. **e**, Changes in PETHs between the PRE and POST periods. Details follow those of Figure 3. Baseline firing rates of both INC- and DEC-type neurons decreased significantly, while movement-related activity was affected only weakly. No significant changes were observed in the peak amplitude of the PETHs in INC-type neurons, the onset timing of movement-related modulations, and modulation duration, except for the trough amplitude in DEC-type neurons.
